# Identity, ontogeny, and age-related changes in splenic white pulp macrophages in mouse and human spleen

**DOI:** 10.64898/2026.04.02.716095

**Authors:** Kalyn R. Thayer, Maxwell J. Schleck, Yuliana V. Sokolenko, Radmila A. Nafikova, Zhan Yu, Ziyan Lu, Karolina J. Senkow, Elsie G. Bunyan, William T. Plodzeen, Constance E. Runyan, Rogan A. Grant, Suchitra Swaminathan, Duc Phan, Hiam Abdala-Valencia, Chitaru Kurihara, Ankit Bharat, Anthony D. Yang, Ryan P. Merkow, Stephanie C. Eisenbarth, Natania S. Field, Samuel E. Weinberg, Mary Carns, Harris Perlman, G.R. Scott Budinger, Alexander V. Misharin

**Author notes:** Corresponding author: Alexander Misharin.

## Abstract

The spleen contains diverse macrophage subsets that remove aged erythrocytes, prevent the dissemination of circulating pathogens, and shape the adaptive immune response^1–3^. The mouse spleen hosts red pulp macrophages (RPM), marginal zone macrophages (MZM), marginal zone metallophilic macrophages (MMM), and tingible body macrophages (TBM). However, their transcriptomic identity, ontogeny, and dynamics during aging are unknown. Furthermore, it is not known whether homologous populations of macrophages exist in the human spleen. We find that in mice, MZM and MMM are tissue-resident macrophages that maintain their population via local proliferation, while TBM are slowly replaced by circulating monocytes. Lineage tracing shows that MMM maintain the MZM pool, and that after MMM depletion, circulating monocytes restore MMM. We show that a decrease in MMM abundance in aging precedes changes in other cellular populations and splenic niches. In human spleen, we identify TBM and perifollicular zone macrophages (PFZM) as a single macrophage population homologous to MMM and MZM in mice. We show that in both mouse and human TBM become more abundant during aging. Our results suggest age-related changes in the splenic microenvironment drive changes in tissue-resident splenic macrophage populations with potential importance for the loss of immunologic function in older individuals.

## MAIN

The spleen is a central lymphoid organ that plays a key role in the innate and adaptive response to infection. For example, patients who undergo splenectomy have a lifetime risk for lethal infections, including sepsis and meningitis caused by pathogens such as *S. pneumoniae, N. meningitidis, H. influenza*^4,5^. This is in part thought to result from a failure to clear encapsulated bacteria that have entered the systemic circulation. Spleen exhibits age-related morphological changes^6,7^, and the phagocytic function of the spleen, mediated by macrophages, declines during normal aging^8,9^, likely contributing to the disproportionate morbidity and mortality associated with common infections in the elderly. However, our understanding of the composition, ontogeny, localization, and age-related changes of splenic macrophage populations, particularly those in the white pulp, is limited.

Red pulp macrophages (RPM) originate from the fetal erythro-myeloid progenitors and fetal liver monocytes and persist with minimal input from the bone marrow monocytes over the lifespan of the mouse during normal aging^10^. RPM are abundant in the spleen and are responsible for the removal of aged erythrocytes and recycle iron^11,12^. The spleen harbors several other mouse macrophage populations, including marginal zone macrophages (MZM), marginal zone metallophilic macrophages (MMM), and tingible body macrophages (TBM)^1^. While these populations of macrophages are relatively rare, genetic and pharmacologic depletion strategies suggest a key role for one or more of these macrophage subsets in controlling bacterial sepsis^13^, viral infections^14^, shaping effective T and B cell responses^15^, preventing autoimmunity^16,17^, and inducing tolerance^18^. However, a lack of validated markers to identify these cells has precluded an understanding of their ontogeny and function and their changes, if any, with advancing age. Furthermore, after more than 40 years since their discovery in rodents^19–24^, it is not known whether similar macrophage populations are present in the human spleen or whether age-related changes in the human spleen are similar to those in the mouse. Here, we combine single-cell genomics, spatial transcriptomics, and lineage tracing to definitively identify the transcriptome, spatial organization, and ontogeny of splenic macrophage populations in the murine and human spleen over the lifespan. Despite important differences between macrophage populations in the murine and human spleen, we identify shared changes in the abundance and spatial localization of splenic macrophages with advancing age in mice and humans that might contribute to the decline in immune function in old animals and the elderly. Single-cell and spatial data can be explored using interactive data browsers at: https://sqlifts.fsm.northwestern.edu/public/Thayer_2026/.

## RESULTS

### Transcriptomic identity of splenic macrophages

CX3CR1, the fractalkine receptor, is highly expressed on monocytes and many macrophage populations throughout the body^10^. We hypothesized that large GFP-positive cells observed in the white pulp and marginal zone of transgenic *Cx3cr1*^gfp/wt^ reporter mice^25^ represent TBM, MMM, and MZM. Immunofluorescent microscopy confirmed that GFP-positive cells in the marginal zone of *Cx3cr*^gfp/wt^ mice were positive for CD169 and MARCO, which were previously identified as specific markers for MMM^26^ and MZM^27^, respectively (**Figure 1a**). We flow cytometry sorted GFP^+^MERTK^+^ cells from the spleen of *Cx3cr*^gfp/wt^ mice (**Figure 1b**). Cytospin analysis identified large cells with a typical macrophage morphology and occasional evidence of recent phagocytosis (**Figure 1c**).

**Figure 1:**
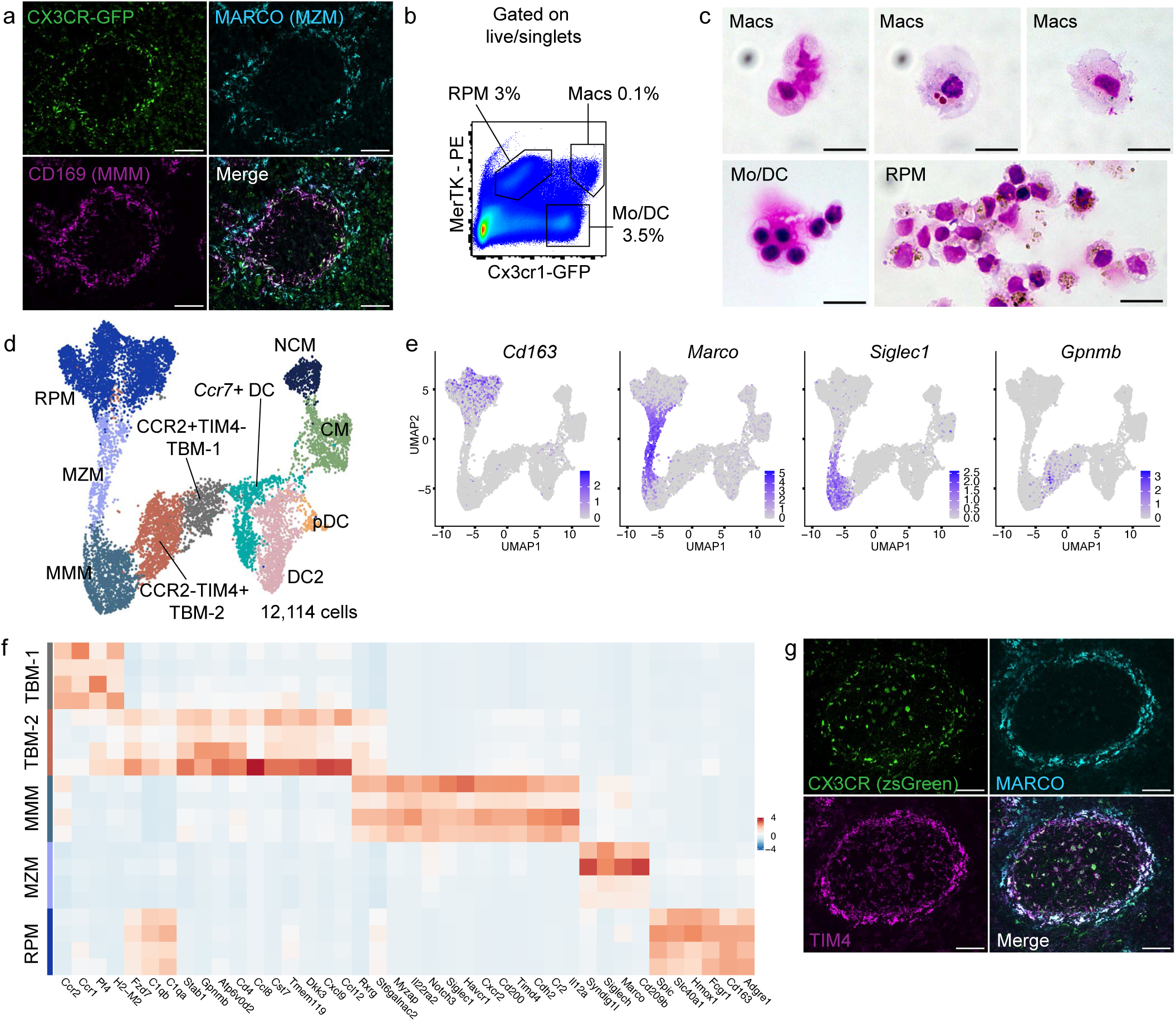
Identification and characterization of macrophages in mouse spleen. a. Immunofluorescence showing GFP fluorescence (green, top left), MARCO staining for MZM (cyan, top right), CD169 staining for MMM (magenta, bottom left), and merge of all three channels (bottom right) from *Cx3cr1^gfp/wt^* mouse spleen. Scale bars 200 μm. b. Gating strategy for flow cytometry cell sorting used to isolate cells for cytospin analysis, starting with live singlets. c. Representative cytospin images. Scale bars 10 μm. d. UMAP plot showing integrated analysis of 12,114 myeloid cells from 3–4-month-old *Cx3cr1^gfp/wt^* mice, 2 male and 2 female. e. Expression of *Cd163*, *Marco*, *Siglec1*, and *Gpnmb* in the myeloid cells on the UMAP in **d**. f. Heatmap showing expression of selected genes within splenic macrophage clusters. Each row represents a single mouse. g. Immunofluorescent microscopy showing endogenous ZsGreen signal (green, top left), MARCO staining for MZM (cyan, top right), Tim-4 staining of MMM, MZM, and TBM (magenta; bottom left), and merge (bottom right) from *Cx3cr1^ERCre^* x ZsGreen mouse spleen. Scale bars 150 μm.

To understand the transcriptional identity of these cells, we performed CITE-seq profiling of flow cytometry-sorted GFP+ cells from four *Cx3cr*^gfp/wt^ mice using a targeted antibody panel (see **Methods**). We resolved clusters corresponding to monocytes, dendritic cells, neutrophils, natural killer cells, and endothelial cells (**Figure 1d**, **Figure S1a, b, c, d, Supplemental Table 1**). Within cells expressing pan-macrophage markers *Mertk, Csf1r, and Mafb*^28^ we identified five subclusters. We then labeled three of these clusters as RPM, MZM, and MMM based on specific markers reported in the literature (**Figure 1e**). These include genes involved in heme and iron metabolism for RPM (*Cd163, Slc40a1, Hmox1*, *Spic*); scavenger receptors (*Cd209b* and *Marco*) for MZM; and sialoadhesin (*Siglec1,* which encodes CD169) for MMM^3,21^ (**Figure 1f**). In addition, we resolved a small subcluster of hemoglobin-containing RPM, possibly representing erythroblastic island RPM (**Figure S1c**). The remaining two macrophage clusters expressed marker genes that were not previously associated with a specific subset of splenic macrophages. These clusters expressed *Pf4, Stab1, Cst7, Cd4,* high levels of cytokines *Cxcl9, Ccl8, Ccl12*, complement component C1q (*C1qa, C1qb, C1qc*), genes related to Wnt-signaling (*Dkk3, Fzd7*), *St6galnac2*, a sialyltransferase, which inhibits the proliferation of B cells and limits the size of spleen germinal centers^29^, and *Gpnmb,* a suppressor of T cell activation^30^ (**Figure 1e, f, Figure S1c, d**). Based on the expression of these markers, we reasoned that these cells likely represent the T and B cell zone-associated TBM. The first TBM cluster, which was characterized by expression of *Ccr1, Ccr2*, and relatively higher levels of MHC II genes, was labeled as CCR2^+^TIM4^-^ TBM (TBM-1). The second TBM cluster lacked expression of *Ccr2*, but expressed *Rxrg*, a retinoid X receptor gamma, *Timd4*, a phosphatidylserine receptor involved in efferocytosis and a modulator of T cell activation^31,32^, and *Atp6v0d2*, which is involved in autophagosome-lysosome function. We labeled this cluster CCR2^-^TIM4^+^ TBM (TBM-2). As expected, MMM, MZM, and TBM uniformly expressed *Cx3cr1* (**Figure S1d**).

Consistent with a role in regulating T and B cell responses, MMM were characterized by expression of genes encoding complement receptor 2 (*Cr2*), TIM1 (*Havcr1*), TIM4 (*Timd4*), IL-22 receptor antagonist (*Il22ra2*), IL-12 subunit alpha (*Il12a*), and a modulator of T cell responses CD200 (*Cd200*), among other genes. Supporting the proposed role of MMM as a cellular barrier that prevents free flow of antigens into the white pulp^33,34^ we found that MMM expressed cadherin *Cdh2* and myocardial *zonula adherens* protein *Myzap*. MZM expressed genes related to phagocytosis, including *Cd209b*, *Cd209d*, *Marco*, *Siglech*, and unlike MMM, genes encoding complement component C1q (**Figure 1f, Figure S1d**).

We assessed expression of protein markers in transcriptomic clusters using CITE-seq (**Figure S1e, f**). F4/80 and CD64 were specific markers of RPM and were not detected on TBM, MMM, and MZM. While *Mertk* expression was uniformly high across TBM, MMM, and MZM, MERTK protein was detected only on CCR2^-^TIM4^+^ TBM-2 and RPM. Detection of CD11b, CD11c, CX3CR1, and I-A/I-E (MHC II) proteins matched mRNA expression patterns and did not further resolve TBM, MMM, and MZM subsets. CD169 (*Siglec1*) was specific to MMM, but its expression was low, limiting its utility for flow cytometry. TIM4 and CD4 were useful to distinguish MMM, TBM and MZM. TIM4 was expressed on 100% of MMM and CCR2^-^TIM4^+^ TBM-2 and the majority of MZM, while CD4 was highly expressed on both subsets of TBM. We used flow cytometry and immunofluorescent microscopy (IFM) to credential TIM4 as a marker to identify MMM, MZM, and the majority of TBM (**Figure 1g**, **Figure S1g**).

We asked whether the expression of specific transcription factors distinguishes splenic macrophage populations (**Figure S1h**). All splenic macrophage subsets expressed *Mafb* and *Zeb2*^35^. As expected, *Spic* and *Pparg* were RPM-specific transcription factors^11,12,28,36^. RPM express *Cebpb*, while TBM, MMM, and MZM express *Cebpa*. Investigators reported that loss of *Nr1h3*, which encodes liver X receptor alpha, a nuclear receptor responsive to oxysterol signaling, led to ablation of MMM and MZM, but not RPM^37,38^. Interestingly, we found that *Nr1h3* was expressed in all splenic macrophage subsets, including RPM. In contrast, the expression of *Rarb*, which encodes retinoid acid receptor beta, was specific to TBM, MMM, and MZM. Expression of *Rxrg*, a retinoid X receptor gamma, was restricted to CCR2^-^TIM4^+^ TBM-2 and MMM. Finally, the expression of *Atf5*, a transcription factor from the CREB family, was restricted to MMM. We did not find a transcription factor uniquely expressed in MZM. Together, these data suggest that splenic white pulp macrophages, particularly MMM and MZM, represent a continuum of states, rather than distinct cell types.

### Splenic MMM, MZM, and TBM are long-living tissue-resident macrophages

While RPM have been shown to be tissue-resident macrophages^10^, it is unknown if MMM, MZM, and TBM rely on local proliferation or continual influx of monocytes for their maintenance at steady state. Lineage tracing in whole body-irradiated bone marrow chimeras demonstrated that splenic MMM and MZM can be repopulated from bone marrow-derived precursors^39^ and expression of *Ccr2* on splenic CCR2^+^TIM4^-^ TBM-1 (**Figure S1e**) suggested that these cells may originate from classical monocytes.

To determine the ontogeny of TBM, MMM, and MZM in adult mice under normal conditions, we performed a lineage tracing experiment using *Cx3cr1*^ERCre^ mice^10^ crossed to ZsGreen reporter mice^40^. We administered a single dose of tamoxifen to label all *Cx3cr1*-expressing cells, including TBM, MMM, and MZM, and assessed the fraction of ZsGreen-positive cells among Tim-4-positive macrophages over a period of 6 months (**Figure 2a, Figure S2a**). Two weeks after tamoxifen treatment, over 90% of Tim4-positive macrophages in both male and female mice were ZsGreen-positive (**Figure 2b**). As circulating classical monocytes are replaced by unlabeled cells from the bone marrow within 2 weeks in this system, the percentage of splenic macrophages without the ZsGreen label indicates replacement by circulating monocytes. Up to 3 months after the tamoxifen pulse, the fraction of ZsGreen-positive macrophages did not significantly decline in male mice. In female mice, the fraction of ZsGreen-positive Tim-4 macrophages did not differ from the baseline at 2 months, but exhibited a small statistically significant decline at 3 months after tamoxifen pulse. Six months after tamoxifen pulse, 73.6% and 79.0% of Tim-4 positive macrophages remained zsGreen positive in males and females, respectively. These results suggest some of the Tim-4 macrophages in the spleen are slowly replaced with monocyte-derived cells. In agreement with our CITE-seq data, which demonstrated that some RPM had detectable *Cx3cr1* expression (**Figure S1e**), we found that a small fraction of RPM was labeled at 2 weeks (5% for male and 10% for female mice) and remained relatively stable for 6 months (**Figure S2b**).

**Figure 2:**
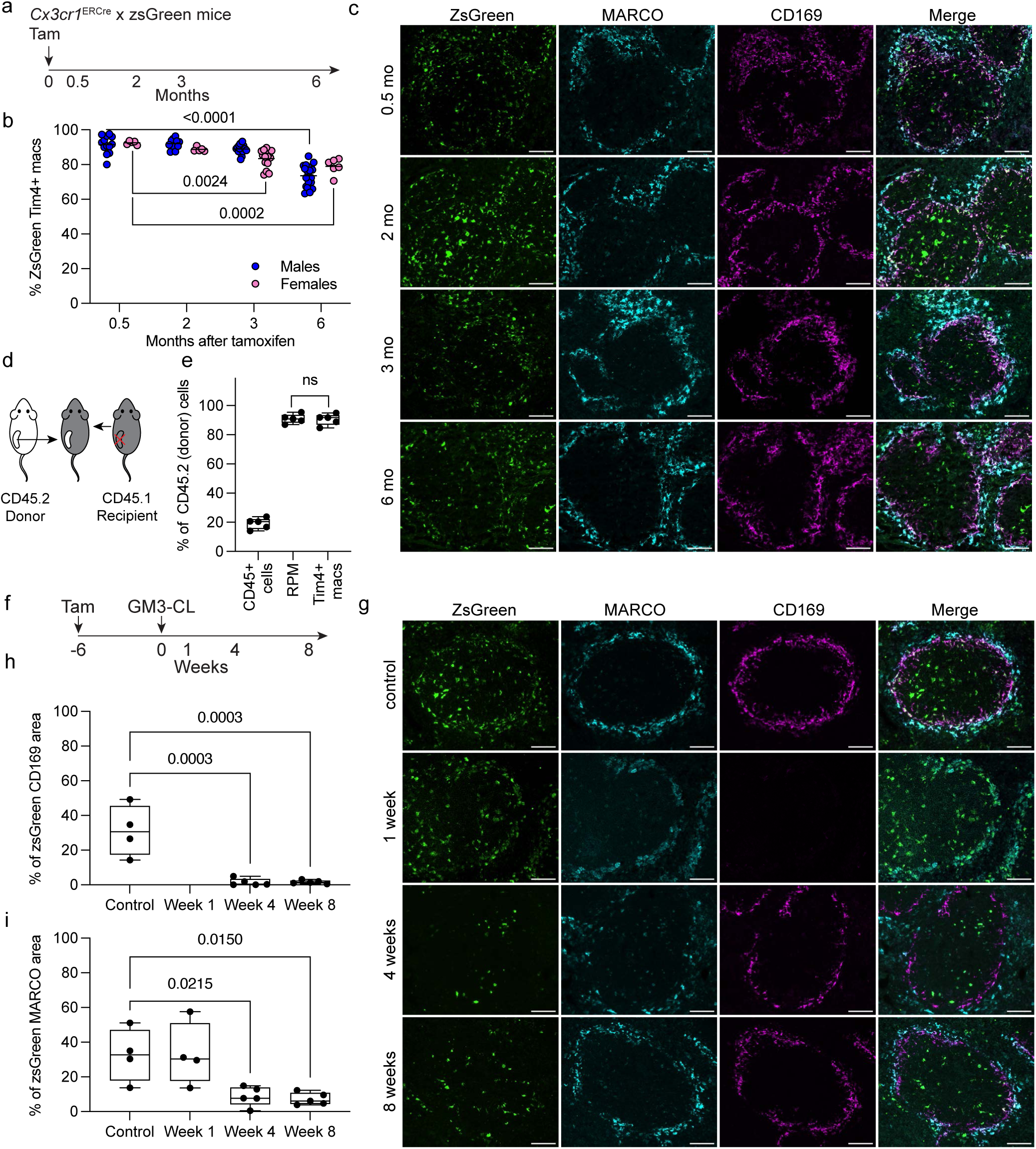
MMM and MZM are tissue-resident macrophages, MMM serve as a precursor population to MZM, and TBM have a dual ontogeny. **a.** Schematic of the lineage-tracing experiment. Tam: tamoxifen. b. Scatterplots showing percentage of ZsGreen-positive Tim-4+ macrophages in *Cx3cr1^ERCre^* x ZsGreen mice after tamoxifen treatment. Number of mice (from left to right): Female: 5, 5, 15, 6. Male: 13, 11, 17, 20. Results were compiled from 4 independent experiments. Significance was determined using one-way ANOVA, using a non-parametric Kruskal-Wallis test with Dunn’s multiple comparison test. The exact adjusted p-value is shown. c. Representative immunofluorescence microphotographs illustrating endogenous ZsGreen fluorescence (green), MARCO staining for MZM (cyan), CD169 staining for MMM (magenta), and merge of all three channels in the spleen of lineage-tracing *Cx3cr1^ERCre^*x ZsGreen mouse after tamoxifen treatment. Scale bars 200 μm. d. Schematic of the spleen transplant experimental design. e. Bar plot showing percentage of donor-derived total immune cells (CD45^+^ cells), RPM, and Tim-4+ macrophages (Tim-4 macs) 3 months post-transplant. Data from 2 experiments, a total of 5 mice, male mice were used for both donors and recipients Kruskal-Wallis test followed by Dunn’s post-hoc testing was used to identify statistically significant differences between groups: adjusted p-value is 0.019 for CD45^+^ cells vs. RPM comparison, 0.029 for CD45^+^ cells vs. Tim-4^+^ macrophages, and >0.99 for RPM vs. Tim-4^+^ macrophages. ns: not significant. f. Schematic of the MMM depletion and repopulation experiment. Tam: tamoxifen. GM3-CL: GM3-modified clodronate-loaded liposomes. g. Representative immunofluorescence microphotographs illustrating endogenous ZsGreen fluorescence (green), MARCO staining for MZM (cyan), CD169 staining for MMM (magenta), and merge of all three channels in the spleen of lineage-tracing *Cx3cr1^ERCre^*x ZsGreen mouse after treatment with GM3-CL. Scale bars 200 μm. h. Box plot illustrating the change in percentage of ZsGreen/CD169 double-positive area after GM3-CL treatment. One-way ANOVA followed by Dunnett’s multiple comparison test was used to identify statistically significant differences between all groups. i. Box plot illustrating the change in percentage of ZsGreen/MARCO double-positive area after GM3-CL treatment. One-way ANOVA followed by Dunnett’s multiple comparison test was used to identify statistically significant differences between all groups.

We confirmed these findings via immunofluorescent microscopy, using CD169 and MARCO to identify MMM and MZM, respectively (**Figure 2c**). Six months after the tamoxifen pulse, the majority of MMM and MZM remained ZsGreen-positive, forming a continuous ring around white pulp. We also observed large ZsGreen-positive, CD169-negative cells in the white pulp follicles up to 6 months after tamoxifen treatment, which are likely TBM. In the absence of a unique protein marker expressed in all TBM, we could not determine their relative rate of replacement using immunofluorescence microscopy.

As an orthogonal approach to interrogate splenic macrophage ontogeny, we performed spleen transplants using CD45.1 mice as recipients and CD45.2 mice as donors (**Figure 2d**). After 3 months, mice were harvested, and macrophage and lymphocyte populations in the spleen were assessed using flow cytometry. As expected, we observed significant replacement of donor CD45^+^ immune cells with recipient cells in the spleen (**Figure 2e**). In contrast, 91.0% of RPM and 90.4% of Tim-4 macrophages were still of donor origin (**Figure 2e**). Together, these experiments show that MMM and MZM are tissue-resident macrophages, while a significant fraction of splenic TBM are replaced by circulating monocytes over the period of several months. This differs from lymph nodes, where several recent reports show that lymph node TBM are derived from lymph node-resident precursors and not circulating monocytes^41,42^.

### MMM maintain MZM pool

Having established that MMM and MZM are tissue-resident cells, we interrogated the relationship between these subsets. As MMM and MZM lie in close proximity to one another and share transcriptomic signatures, we considered the possibility that one population may differentiate into the other. To test this, we treated *Cx3cr1*^ERCre^ x ZsGreen reporter mice with tamoxifen to label all MMM, MZM, and TBM. Six weeks later, we treated these mice intravenously with ganglioside GM3-modified clodronate-loaded liposomes (GM3-clo-lip), which specifically deplete MMM^43^. We then harvested spleens 1, 4, and 8 weeks after MMM depletion (**Figure 2f**). If newly emerging MMM are reconstituted from circulating monocytes, they will be zsGreen-negative, while if they are reconstituted by MZM or TBM, they will be zsGreen-positive.

While the overall histology of the spleen was largely unaffected by treatment with GM3-clo-lip (**Figure S2c**), we observed that MMM were almost entirely absent from the spleen one week after the treatment (**Figure 2g,h**). At the 4-week timepoint, ZsGreen-negative MMM started to reappear and form incomplete rings around the marginal zone. After 8-weeks, the population of zsGreen-negative CD169-positive MMM formed nearly complete rings around white pulp (**Figure 2g,h**). The percentage of ZsGreen-positive MARCO-positive MZM was unchanged 1 week after GM3-clo-lip treatment, but declined 4 and 8 weeks later (**Figure 2g,i**). Thus, our data indicate that after depletion MMM are replaced by monocytes, rather than by the trans-differentiation of TBM or MZM. Moreover, consistent with scRNA-seq data, this finding suggests that, while the MZM pool is relatively stable, it is maintained through the trans-differentiation of MMM.

### Age-associated changes in MMM abundance precede changes in splenic architecture

While age-related changes in mouse splenic architecture, composition, and function have been reported^44,45^, systematic evaluation of such changes is lacking. Therefore, we evaluated differences in spleen architecture with age. We quantified the relative contributions of white pulp/marginal zone and red pulp to the total tissue area in young, old, and very-old C57BL/6 male mice. We found that an age-associated loss of the white pulp/marginal zone was only apparent in very old mice (57.2%, 58.7%, 35.2% in young, old, and very old mice, respectively), compensated by a reciprocal increase in red pulp area (**Figure S3a, b**).

To gain a greater understanding of the age-associated changes in the architecture and cellular composition of the spleen, we performed single-cell spatial transcriptomic analysis. First, we assembled a scRNA-seq spleen atlas using our mouse data (**Figure S1a**) and two previously published scRNA-seq datasets^46,47^ (**Figure S3c-e**), which we used to design a custom gene panel for spatial transcriptomics (**Supplemental Table 2**). In spatial data, we resolved 29 clusters, including clusters corresponding to RPM, MZM, MMM, and TBM, and 11 niches, which we hierarchically annotated and harmonized using the mouse single-cell atlas (**Figure 3a–d, Figure S3f–i, Supplemental Tables 3 and 4**). We found that macrophages identified as MZM by their transcriptome were interspersed among MMM and did not form a distinct layer (**Figure 3b**). This observation further supports our scRNA-seq and lineage tracing data showing that that MMM and MZM represent a continuum of cell states. This finding differs significantly from the traditional understanding of marginal zone architecture as static and distinct compartments, which relied on IFM studies using anti-CD209b or MARCO antibodies.

**Figure 3:**
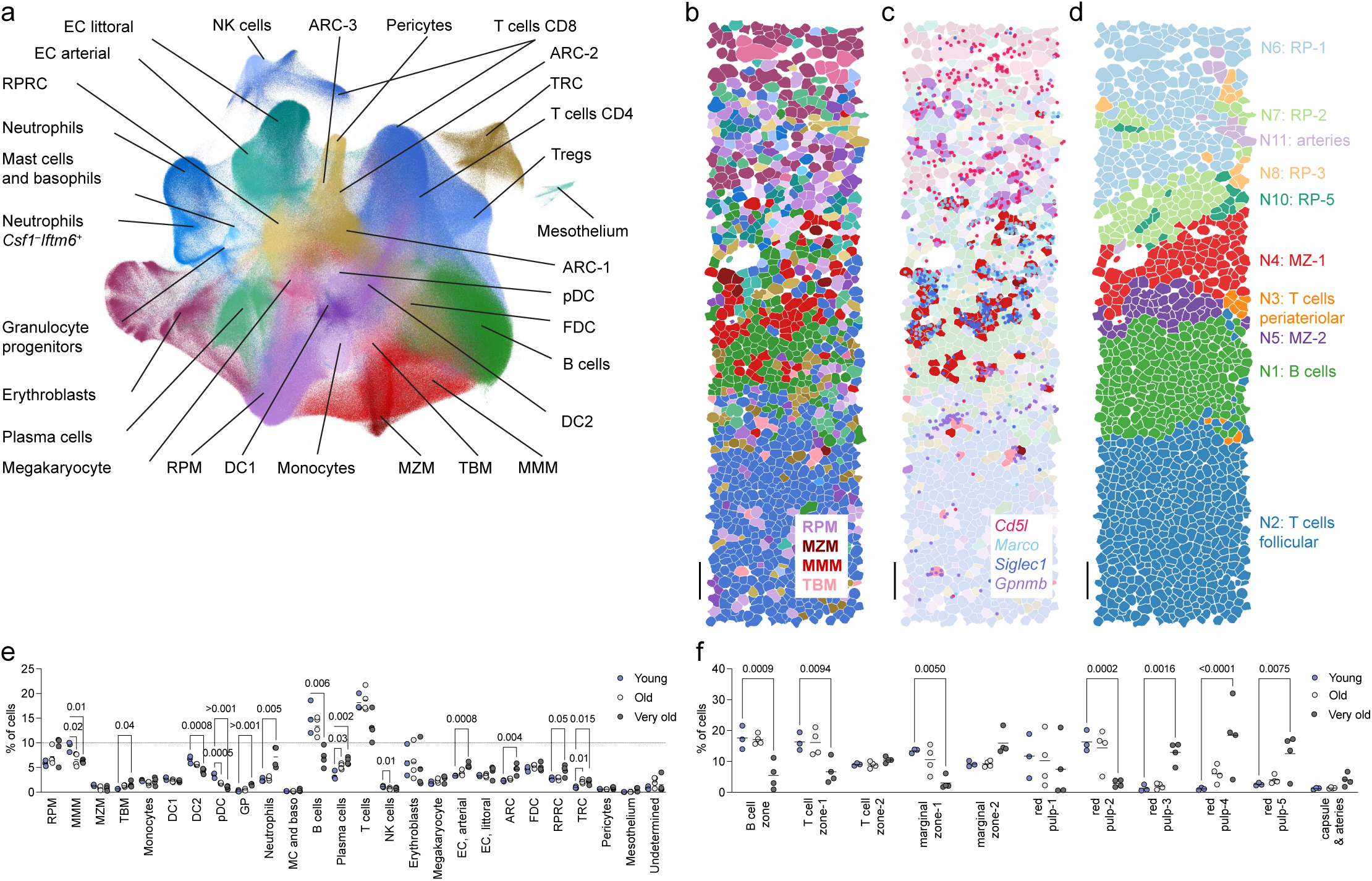
Changes in the aged mouse spleen are independent of a young circulation and include a decline in MMM and an increase in TBM, both of which remain tissue-resident cells. **a.** UMAP plot showing integrated analysis of cells identified via spatial transcriptomics from young adult (n=3, 5-month-old), old (n=4, 22-month-old), and very old (n=4, 32–33-month-old) C57BL/6 male mice. Level 3 clusters are shown. b. Spatial projection of cell types (level 3) resolved via spatial transcriptomics in the young mouse. Color palette is the same as **a**. Scale bar is 25 μm. c. Spatial projection of cell types (level 3) and macrophage-specific marker genes resolved via spatial transcriptomics in the young mouse. Same region as in panel **b**. Color palette is the same as in panels **a** and **b**. Colored dots indicate specific transcripts. Scale bar is 25 μm. d. Spatial projection of niches resolved via spatial transcriptomics. Same region as in panel **b**. Scale bar is 25 μm. e. Scatter plot illustrating changes in relative abundance of splenic cell type (level 3) between young, old, and very old mice. Horizontal bars indicate group means. Two-way ANOVA, followed by Dunnett’s multiple comparison test was used to identify statistically significant differences between all groups. f. Scatter plot illustrating changes in relative abundance of splenic niches between young, old, and very old mice. Horizontal bars indicate group means. Two-way ANOVA, followed by Dunnett’s multiple comparison test was used to identify statistically significant differences between all groups.

Splenic macrophages are integrated with the extracellular matrix and are often fragmented during tissue processing, leading to underrepresentation in tissue composition assessments via flow cytometry^48–50^. Hence, we leveraged our spatial transcriptomics data to determine the relative abundance of macrophages, as well as all other cell types, in the spleen and their changes with age. Surprisingly, we found that in young adult mice, MMM were the most abundant macrophage subset (9.2% of all cells), followed by RPM (5.8%), MZM (1.4%), and then TBM (0.66%) (**Figure 3e**). Of note, current cell segmentation algorithms likely systematically overestimate the abundance of splenic macrophages (**Figure 3c**). In old mice, the relative abundance of MMM declined (6.6%), while the abundance of other macrophage populations did not change (**Figure 3e**). In addition to the decline in MMM abundance, pDCs and NK cells were decreased in spleens from old compared to young adult mice, while plasma cells and TRC were increased (**Figure 3e**). Thus, an increase in MMM abundance was the earliest change in splenic macrophages we observed in the aging spleen.

In contrast to the small changes we observed in old compared to young adult mice, the relative abundance of many cell types in the spleen changed substantially in very old mice when compared to young mice. In very old mice, the most abundant macrophage was RPM (8.8%), followed by MMM (6.6%), TBM (1.5%), and MZM (0.8%). In spleens from very old compared to young adult mice, there was a decrease in the relative abundance of DC2, pDC, and B cells and an increase in stromal cell populations of T cell zone reticular cells (TRC), red pulp reticular cells (RPRC), adventitial reticular cells (ARC), and arterial endothelial cells, as well as granulocyte progenitors, neutrophils, and plasma cells (**Figure 3e**). This increase in granulocyte progenitors and neutrophils in very old mice suggests increased extramedullary hematopoiesis with aging.

Niche analysis resolved 11 niches (**Figure 3d; Figure S3h, i, k, l**). We resolved the B cell zone niche (N1) and two T cell niches: follicular (N2) and periarteriolar (N3), enriched for DC2 and adventitial reticular cell subsets (ARC-1, ARC-2, and ARC-3) (**Figure 3d; Figure S3h,i, k, l**). Within red pulp, which is generally considered homogenous, we resolved five distinct niches: red pulp niche-5 (N10) was formed by littoral endothelial and arterial cells and red pulp reticular cells. Red pulp niche-4 (N9) was characterized by clusters of plasma cells adjacent to white pulp follicles, while red pulp niche-3 (N8) was characterized by clusters of *Mpo+* granulocyte progenitors, immature *Csf1-Iftim6+* and mature *Csf1+* neutrophils. Unlike the “deep” red pulp niche-1 (N6), which was enriched for erythroblasts, red pulp niche-2 (N7) was immediately adjacent to the marginal zone and contained diverse populations of immune cells.

We focused on the niches formed by TBM, MMM, and MZM. Within the marginal zone, we resolved inner (marginal zone-1, N4) and outer (marginal zone-2, N5) niches. While both niches contained an equal fraction of MMM, the inner zone was characterized by the presence of CD4 T cells, *Madcam1+* FDC, and *Gpihbp1*+ arterial endothelial cells, while the outer MMM zone was enriched for B cells, CD8 T cells, DC1, and MZM. As expected, TBM were present in T and B cell zones, as well as in the inner marginal zone niche, and were frequently positioned next to TRC and FDC.

In comparison to young mice, very old mice had significant changes in multiple niches, including a decrease in B and T cell zones, a loss of the outer layer of the marginal zone, and a corresponding increase in the inner layer of the marginal zone (**Figure 3e**). Red pulp niches also differed in very old compared to young adult mice and were characterized by an increase in plasma cell-dense regions and areas of granulocytopoiesis, and a decrease in red pulp adjacent to the marginal zone. In contrast, niche proportions were not different between young adult and old mice.

### Identification of MMM and TBM homologs in human spleen

Surprisingly little is known about macrophage subsets in the human spleen. Human splenic architecture differs substantially from that of the mouse spleen in ways that likely impact splenic macrophage phenotype^1,34,51^; notably, the human spleen lacks a marginal zone sinus lined with macrophages. Instead, it contains a plexus of small arterioles and capillaries that end in a completely open circulation; many of these capillaries are sheathed capillaries containing specialized endothelial cells, stromal cells, and macrophages. Previous immunohistochemistry (IHC) studies on a limited number of human spleens have identified CD169^+^ macrophages as part of the sheathed capillary structure, though only in the perifollicular zone^52^. Additionally, TBM have not been directly studied in human spleen, and no markers have been found to identify them via histology analysis.

To characterize macrophage populations in human spleen, we obtained spleens from 6 patients undergoing splenectomy for non-spleen-related diseases (primarily pancreatic cancer) and 6 organ donors (**Supplemental Table 5**). Since markers for human homologs of TBM, MMM, and MZM were unknown, we performed broad enrichment for myeloid cells using flow sorting followed by scRNA-seq (**Figure S4a**). We resolved clusters corresponding to macrophages, monocytes, subsets of dendritic cells, neutrophils, natural killer cells, T and B lymphocytes, endothelial and stromal cells (**Figure 4a**; **Figure S4b**, **Supplemental Table 6**). We identified two macrophage clusters by expression of known macrophage markers, including *APOE, APOC1, MERTK, AXL, CSF1R,* and *C1QA* (**Figure 4a,b, Figure S4c**). One macrophage cluster was characterized by expression of *HMOX1, CD5L, CD163, VCAM1*, *SLC40A1* (ferroportin gene), and *FCGR3A* (CD16) and matched RPM. The second macrophage cluster was characterized by expression of lipid transfer and metabolism genes such as *LPL*, *PLA2G2D, PTGDS, PLTP,* and *SCARB1* (scavenger receptor class B), *TIMD4*, metalloproteinases *MMP9* and *ADAMDEC1*, *ITGAM* (CD11B), *MSC* (musculin, transcriptional repressor capable of binding an E-box element), *IL2RA* (CD25), *NOTCH3*, and *GAS6*. Notably, this cluster showed very low or undetectable levels of canonical macrophage markers, including *MRC1* (CD206)*, MSR1* (CD204), and *MARCO* (MARCO) (**Figure 4b**).

**Figure 4:**
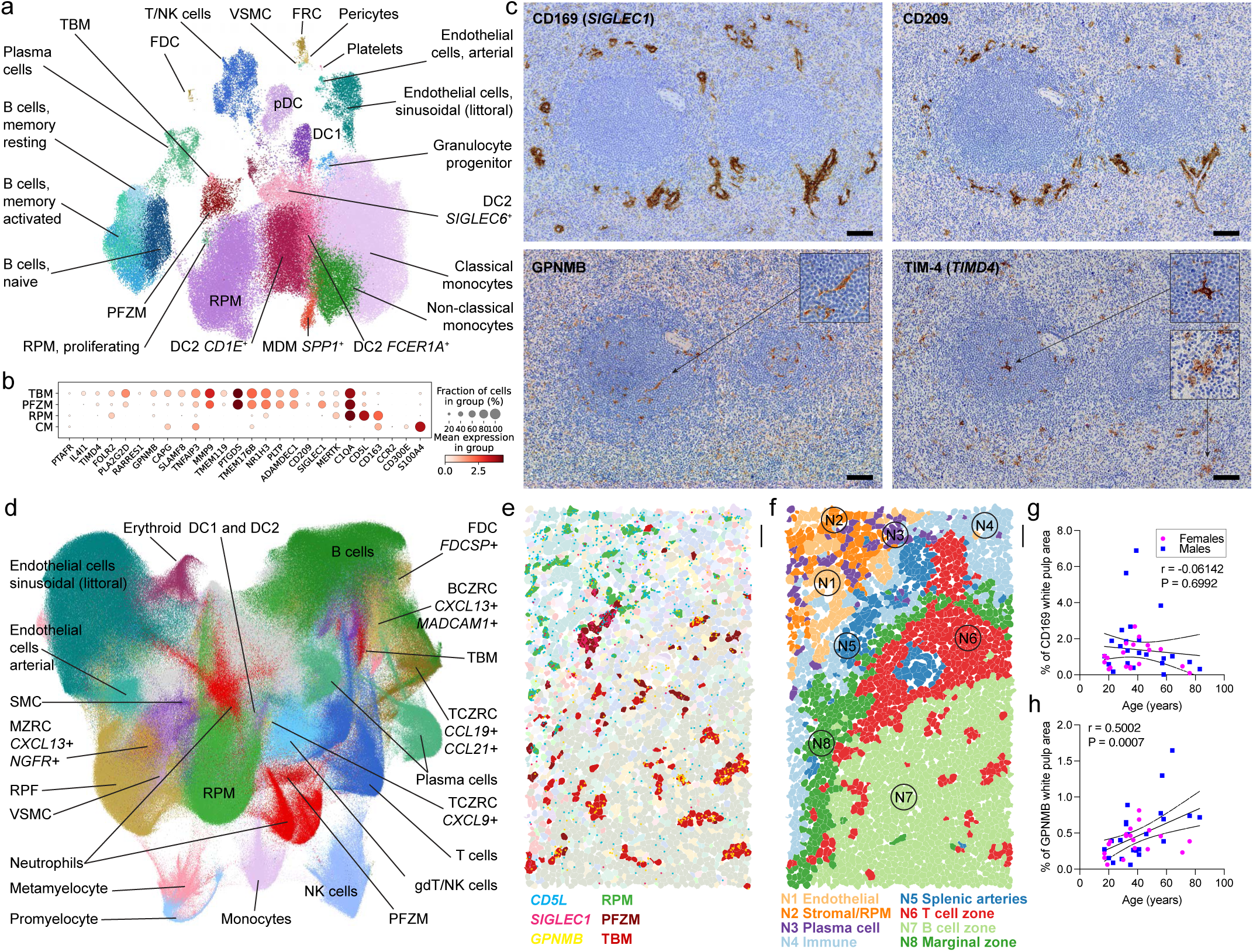
Identification of white pulp macrophages and their age-related changes in human spleen. **a.** UMAP plot showing integrated scRNA-seq analysis of 130,708 cells from 12 human spleen samples. TBM: tingible body macrophages; FDC: follicular dendritic cells; VSMC: vascular smooth muscle cells; RFC: follicular reticular cells; DC1: dendritic cells type 1; DC2: dendritic cells type 2; pDC: plasmacytoid dendritic cells; MDM: monocyte-derived macrophages; RPM: red pulp macrophages, PFZM: perifollicular zone macrophages. b. Dot plot showing expression of select genes in tingible body macrophages (TBM), perifollicular zone macrophages (PFZM), red pulp macrophages (RPM), and classical monocytes (CM). c. Representative immunohistochemistry for CD169, CD209, GPNMB, and TIM-4 on adjacent sections from the same human spleen with hematoxylin counterstaining. Scale bar is 100 μm. d. UMAP plot showing integrated single-cell spatial transcriptomic analysis of 2,428,938 cells from 8 human spleens. Level 3 cluster annotations are shown. BCZRC: B cell zone reticular cells; TCZRC: T cell zone reticular cells; RPF: red pulp fibroblasts; MZRC: marginal zone reticular cells; SMC: smooth muscle cells. e. Spatial projection of cell types (level 3) and macrophage-specific marker genes in the human spleen resolved via spatial transcriptomics. Color palette is the same as **d**. Scale bar is 25 μm. f. Spatial projection of niches resolved via spatial transcriptomics. Same region as in panel **f**. Scale bar is 25 μm. g. Scatter plot showing correlation between age and percentage of CD169-positive area within the splenic white pulp from 42 subjects. The solid line illustrates the correlation, and the dashed line illustrates the 95% confidence interval. Scatter plot showing correlation between age and percentage of GPNMB-positive area within the splenic white pulp from 42 subjects. The solid line illustrates the correlation, and the dashed line illustrates the 95% confidence interval.

Within this second macrophage cluster, we resolved two subclusters. The first subcluster was characterized by expression of *SIGLEC1* (CD169) and *CD209* (known as DC-SIGN), and matched gene expression profiles of MMM and MZM in the mouse spleen (**Figure 4b; Figure S4c,d**). The second subcluster expressed *ITGAX* (CD11C) and *GPNMB*, matching gene expression profiles of mouse TBM (**Figure 4b; Figure S4c,d**). Both subclusters had variable expression of *TIMD4.* Importantly, all macrophage clusters contained cells from all 12 subjects (**Figure S4e**). IHC for CD169, CD209, GPNMB, and TIM4 confirmed specific location of these macrophages in a larger cohort of 42 human spleen samples (**Supplemental Table 5**). CD169- and CD209-positive macrophages formed rings and tracks located mainly in perifollicular regions and marginal zone (**Figure 4c**). These CD169 cells matched the ring-shaped morphology of previously-reported perifollicular zone macrophages (PFZM)^52,53^. IHC for GPNMB identified large cells, located inside follicles, with long processes and occasional foamy cytoplasm (**Figure 4c**). TIM4 staining was highly variable between samples, but some samples had large TIM4-positive cells inside follicles and marginal zone (**Figure 4c**). Based on their gene expression and location in the white pulp, we annotated the first subcluster of *SIGLEC1^+^* macrophages as PFZM. We interrogated transcription factors and transcriptional regulators expressed in PFZM and TBM (**Figure S4f**). Notably, both *NR1H3* (encodes LXRa) and *NOTCH3* were highly expressed in PFZM and TBM, mirroring mouse MMM and TBM. Other transcription factors shared between mouse and human were *MAF, CEBPA, RXRA*. The only transcription factor that differed between TBM and PFZM was *RELB,* which was highly expressed in TBM only.

To define the spatial niches for PFZM and TBM in the human spleen, we performed single-cell spatial transcriptomics on 8 human spleens from organ donors or patients undergoing splenectomy for pancreatic cancer (**Supplemental Table 5**). Clustering on transcriptomic profiles resolved multiple cell types, including stromal, endothelial, and immune cells, which we hierarchically annotated and harmonized with the human scRNA-seq data (**Figure 4d; Figure S4g; Supplemental Tables 7 and 8**). In addition to RPM (7.7% of cells), we identified clusters of rare cells that matched gene expression profiles of PFZM and TBM (0.35% and 0.32%, respectively, **Figure 4d, Figure S4h**). Importantly, since current cell segmentation and transcript assignment approaches are imperfect, single-cell spatial transcriptomics likely significantly overestimates the number of PFZM and TBM (**Figure 4e**).

Niche analysis resolved 8 niches that matched known anatomical structures in the spleen (**Figure 4f, Figure S4j,k**). Within the red pulp, we resolved an endothelial niche (N1), formed by littoral endothelial cells, a stromal/RPM niche (N2), formed by RPM and red pulp fibroblasts, a plasma cell niche (N3), formed by plasma cells and dendritic cells, and an immune niche (N4), formed by RPM, endothelial cells, and immune cells typically found in circulation. The splenic arteries niche (N5) was formed by vascular smooth muscle cells, arterial endothelial cells, and surrounding immune cells. Within the white pulp niche, we resolved follicular T and B cell niches and the marginal zone niche. T cell niche (N6) was organized around splenic arteries and, in addition to T cells, contained *CCL19^+^ CCL21^+^* T cell zone reticular cells and TBM. B cell zone niche (N7) was characterized by *CXCL13* expression and presence of *FDCSP*^+^ fibroblastic reticular cells. Marginal zone niche (N8), surrounding B and T cell zones, was enriched for B cells and fibroblastic reticular cells, including *CXCL13^+^* and *CXCL13^+^MADCAM1^+^*B cell zone fibroblastic reticular cells. PFZM were enriched in the splenic arteries niche (N5), the adjacent perifollicular T cell zone (N6), and the marginal zone niche (N8). TBM were enriched in the marginal zone niche (N8, 35.0%) and the follicular B cell zone niche (N7, 34.3%).

Interestingly, while in healthy adult humans spleen is generally not considered a primary site of granulopoiesis, in all seven subjects, we detected *MPO^+^ELANE^+^* promyelocytes and DEFA1+ metamyelocytes/band neutrophils, which also expressed *MKI67* and *PCNA*, indicative of ongoing proliferation (**Figure S4g,h**). The majority of promyelocytes were located in the red pulp plasma cell niche (N3) and red pulp immune niche (N4) (51.2% and 34.8%, respectively), while the majority of metamyelocytes were located in the red pulp immune niche (34.8%) (**Figure S4k**).

### TBM are expanded in the human splenic white pulp during aging

We set out to determine if the relative abundance of TBM and PSM change with age in human spleen by quantifying IHC staining of GPNMB and CD169 in the extended cohort of 42 subjects (19 females, 23 males). Since area of the white pulp substantially varies between subjects and does not exhibit correlation with age or sex (**Figure S4k**), we normalized the area of GPNMB- or CD169-positive staining to the total white pulp area for each sample. We found that white pulp TBM increased in relative abundance with age (**Figure 4f**); this correlation was statistically significant for males but not females when split by sex (**Figure S4l**). We found no correlation between age and relative abundance of CD169^+^ PFZM in white pulp (**Figure 4f, Figure S4m**).

## DISCUSSION

This study provides a unified transcriptional, spatial, and lineage-resolved framework for splenic white pulp macrophages, resolving long-standing ambiguity surrounding marginal zone macrophages (MMM/MZM) and tingible body macrophages (TBM). By integrating scRNA-seq, fate mapping, depletion-repopulation experiments, and spatial transcriptomics, we demonstrate that MMM and MZM constitute a continuum of tissue-resident macrophage states maintained locally, with MMM functioning as a progenitor-like population that sustains the MZM pool. In contrast, TBM exhibits mixed ontogeny, with a substantial monocyte-derived component even at steady state. These findings revise classical splenic macrophage hierarchies derived largely from static histology and establish ontogenetic relationships between subsets.

A central insight of this work is that age-associated changes in splenic macrophage composition are initiated early, preceding gross architectural remodeling of the spleen. We identify a selective decline in MMM abundance as the earliest detectable cellular change during aging, occurring before contraction of white pulp or marginal zone niches. Future studies will identify whether this decline is due to increased cell-autonomous macrophage turnover, loss of systemic rejuvenating factors, or age-dependent alterations in the local stromal microenvironment.

By extending our analysis to human spleen, we identify perifollicular zone macrophages (PFZM) as transcriptional and spatial homologs of mouse MMM/MZM, alongside *bona fide* TBM populations within follicles. Despite architectural differences between mouse and human spleen, key transcriptional regulators (including *NR1H3, RXR* family members, and *NOTCH* signaling components) are conserved, suggesting evolutionary preservation of white pulp macrophage programs. Notably, aging in human spleen mirrors mouse findings in a subset-specific manner, with expansion of GPNMB⁺ TBM but relative stability of CD169⁺ PFZM abundance. We found that age-related expansion of GPNMB^+^ TBM was specifically observed in males, but not in females. This is potentially important, as many autoimmune diseases, such as systemic lupus erythematosus, are more prevalent in women. This divergence further suggests that distinct macrophage niches respond differently to aging cues in humans, with potential implications for age-related defects in humoral immunity and apoptotic cell clearance. Future work will determine whether this age-related expansion of TBM plays a protective or maladaptive role in maintaining T and B cell-mediated responses to infection, vaccination, or cancer.

## METHODS

### Human spleen tissue collection

All human research was approved by the Northwestern University Institutional Review Board (STU00214971, STU00215396, STU00212120). Human spleen samples were obtained from patients undergoing splenectomy during elective surgery (primarily for pancreatic cancer resection) or from organ donors. Splenectomy samples (12/45 samples) were obtained via NDRI (protocol RMIA2; STU00214971) or the Surgical Oncology program at Northwestern Medicine (STU00215396). Spleens from patients who had ever undergone chemotherapy or radiation were excluded from the study. Spleens from organ donors (32/45 samples) were obtained via the Lung Transplant program at Northwestern Medicine (STU00212120), all study participants or their surrogates provided informed consent. Age of samples in the complete cohort ranged from 17–83 years, with an average age of 39.8. 32/42 (76.2%) samples were from white patients or tissue donors, and 23/42 (54.8%) were male. Full demographic characteristic of the cohort is reported in **Supplemental Table 5**.

### Animal studies and mouse models

All experimental protocols were approved by the Institutional Animal Care and Use Committee at Northwestern University. All strains were housed at a barrier- and specific pathogen-free facility at the Center for Comparative Medicine at Northwestern University. We used definitions for murine age from the Jackson Laboratory, which defines mature adult/young adult mice as 3–6 months old, middle-aged mice as 10–14 months old, and old mice as 18–24 months of age, and very old mice as >24 months of age. The *Cx3cr1-Cre*^ERT2^ mice^10^, zs-Green reporter mice^40^, and *Cx3cr1*^gfp^ reporter mice^25^ were obtained from Jackson Laboratories (Jax stocks 020940, 007906, and 005582, respectively).

### Oral gavage of tamoxifen

For induction of Cre-ERT2 activity, 10 mg tamoxifen (Sigma, T5648) was dissolved in 100µL corn oil (Sigma, C8267) and administered to the anesthetized mice by oral gavage. Mice were treated once.

### Spleen transplantation

After anesthesia and preparation of the surgery site, donor mouse was given a heparinized saline injection through IV. The thorax was then opened and the right atrium incised to allow blood to exit during perfusion. The entire mouse was then perfused with a total of 15 ml of normal saline through a cannula inserted into the apex of the left ventricle. Then, the abdomen of the donor was opened with a longitudinal incision. Small vessels between the pancreas and the intestine were ligated with 7-0 silk suture. The celiac artery was then isolated, and the hepatic and gastric artery ligated with 7-0 silk suture. The abdominal aorta was ligated and cut just below the celiac artery and also dissected above the celiac artery. This approach resulted in an aortic cuff connected to the splenic artery, which allowed vascular anastomosis of the spleen to the recipient. Following ligation of the bile duct, the portal vein was isolated, and the superior and inferior mesenteric and gastric veins were ligated. The portal vein was intersected closely to the liver. The entire organ package containing the vascular connections, spleen and the pancreas was then removed and stored in ice cold saline for 15 minutes while the recipient was prepared. Recipient mouse was anesthetized and the surgery site was prepared for aseptic surgery. An abdominal midline incision was made and the inferior vena cava and the descending aorta isolated below the renal arteries. The recipient vessels were clamped with vessel clamps and opened with microscissor. The portal vein was anastomosed to the inferior vena cava and the donor aortic cuff was connected with an end-to side anastomosis to the recipient aorta using 10-0 suture. The clamp was then removed to restore blood flow. Before closure of the abdomen, the recipient spleen was removed by ligating all the vessels supply to the spleen with a 7-0 silk suture. The abdomen wall is closed in two separate layers using 5-0 vicryl (inner layer) followed by 5-0 non-absorbable nylon sutures. Ophthalmic ointment was applied to the corneas in any sedation/anesthesia > 20 minutes to prevent corneal drying that can lead to ulceration. Mice received Buprenorphine SR (Covetrus) to control post-operation pain. Mice also received 20mg/kg Meloxicam if they showed signs of breakthrough pain and subcutaneous normal saline if they showed signs of dehydration during the first few days after surgery.

### Pharmacological MMM depletion

To specifically deplete MMM, mice were treated with 100 ul of modified GM3 clodronate-loaded liposomes (standard liposome preparation modified with 3% mol GM3 ganglioside, then size extruded via dialysis through 200 nm filter; https://clodronateliposomes.com/) delivered via single retro-orbital i.v. injection under light isoflurane anesthesia. Control mice received PBS.

### Single cell suspension preparation

#### Mouse

To prepare a single cell suspension for flow cytometry analysis of myeloid cells, cytospin analysis, or CITE-seq, approximately quarter of the whole mouse spleen was infused with 2–2.5 ml of digestion buffer containing 2 mg/ml collagenase I (Roche) and 200 μg/ml DNase (Roche) in PBS (containing Ca and Mg) using a syringe with a 30G needle. Spleen was incubated for 30 min at 37°C with gentle agitation. At 15 minutes into incubation, spleen tissue was gently pipetted using wide bore tip. Following digestion, the spleen was pushed through at a 70 μm cell strainer (Miltenyi SmartStrainers) using a syringe plunger, and the cell strainer was rinsed with 10-20 ml of MACS buffer (Miltenyi). After centrifugation at 400 rcf for 10 min at 4°C, the supernatant was removed, and erythrocytes were lysed by resuspending the cell pellet in 2 ml of PharmLyse buffer (BD) and incubated for 2 min on ice with shaking. Lysis was stopped by adding 20 mL of MACS buffer, followed by centrifugation at 400 rcf for 10 min at 4°C. Cells were resuspended in MACS buffer, and cell concentrations and viability were measured using a Cellometer K2 with Acridine Orange/Propidium Iodide (AO/PI) reagent (Nexcelom).

To prepare a single cell suspension for lymphocyte flow analysis, approximately quarter of the whole mouse spleen was placed on polystyrene weighing dish and mashed gently with a syringe plunger until a homogenous pulp was formed. The mashed spleen pulp was then pushed through at a 70 μm cell strainer (Miltenyi SmartStrainers) using a syringe plunger, and the cell strainer was rinsed with 10–20 ml of MACS buffer (Miltenyi). After centrifugation at 400 rcf for 10 min at 4°C, the supernatant was removed, and erythrocytes were lysed by resuspending the cell pellet in 2 ml of PharmLyse buffer (BD) and incubated for 2 min on ice with shaking. Lysis was stopped by adding 20 mL of MACS buffer, followed by centrifugation at 400 rcf for 10 min at 4°C. Cells were resuspended in MACS buffer, and cell concentrations and viability were measured using a Cellometer K2 with AO/PI reagent.

#### Human

To prepare a single cell suspension, any visible capsular tissue was removed and a roughly 1 cm^3^ piece of spleen was infused with 5 ml of digestion buffer containing 2 mg/ml collagenase I (Roche) and 200 μg/ml DNase (Roche) in PBS (containing Ca and Mg) using a syringe with a 30G needle. Spleen was cut into several smaller pieces using scissors and then incubated for 30 min at 37°C with gentle agitation. Following digestion, the spleen was pushed through at a 70 μm cell strainer (Miltenyi SmartStrainers) using a syringe plunger, and the cell strainer was rinsed with 25 ml of MACS buffer (Miltenyi). After centrifugation at 400 rcf for 10 min at 4°C, the supernatant was removed, and erythrocytes were lysed by resuspending the cell pellet in 15 ml of PharmLyse buffer (BD) and incubated for 2 min on ice with shaking. Lysis was stopped by adding 35mL of MACS buffer, followed by centrifugation at 400 rcf for 10 min at 4°C. Cells were resuspended in MACS buffer, and cell concentrations and viability were measured using a Cellometer K2 with AO/PI reagent. At this point, cell suspension was either frozen for cryopreservation in Bambanker (Bulldog Bio) or stained for cell sorting.

#### Flow cytometry and cell sorting of human splenocytes for scRNAseq

Starting with single cell suspension, (either fresh or thawed cryopreserved cells), 1 - 3 × 10^7^ cells were aliquoted and centrifuged at 400 rcf 5 min at 4°C, then supernatant was removed. Cells were resuspended in 1:50 Fc Block in MACS buffer so that the final concentration of cells was 1 × 10^7^ cells 100µL^−1^. Fluorophore-conjugated antibody cocktail was added in a 1:1 ratio. The following antibodies were used (antigen, clone, fluorophore, manufacturer, catalog no., *final* dilution during incubation, unique antibody identifier): CD14, M5E2, BUV395, BD, 740286, 1:20, AB_3099635; HLA-DR, L243, eFluor450, Invitrogen, 48-9952-42, 1:40, AB_1603291; CD45, HI30, BV510, BioLegend, 304036, 1:20, AB_2561940; CD163, GHI/61, PECy7, BioLegend, 333614, 1:40, AB_2562641; CD169, 7-239, APC, BioLegend, 346008, 1:20, AB_11147948; TIM-4, 9F4, PE, BioLegend, 354004, 1:20, AB_11124105; CD209, 9E9A8, PE, BioLegend, 330106, 1:40, AB_1134052; CD19, BV785, HIB19, BioLegend, 302240, 1:20, AB_11218596; CD3, BV785, OKT3, BioLegend, 317330, 1:20, AB_2563507; GPNMB, PE, HOST5DS, Invitrogen, 1:20, 12-9838-42, AB_2572736; CD15, BV785, SSEA-1, BioLegend, 323044, 1:20, AB_2632921; CD15, BV786, HI98, BD, 563838, 1:20, AB_2738444; CD4, BUV737, SK3, BD, 612748, 1:20, AB_2870079.

After incubation at 4 °C for 30 min in the dark, cells were washed with 3 ml of MACS buffer, then centrifuged at 400 rcf 5 min at 4°C. Supernatant was removed, cells were resuspended at a concentration of 1 × 10^7^ cells ml^−1^ and kept on ice until sorting. SYTOX Green viability dye (Thermo Fisher Scientific) was added to cell suspension 5 minutes prior to sorting sample at a concentration of 1 μl per 1 ml. Samples were mixed gently prior to sorting. Samples were filtered through 30 µm pre-separation filters (Miltenyi Biotec) immediately prior to sorting. Cells were sorted on a BD FACS Aria III SORP instrument using a 100-μm nozzle. Cells were sorted into 300 μl of 2% bovine serum albumin (BSA) in Dulbecco’s phosphate-buffered saline (DPBS) and immediately after sorting pelleted by centrifugation at 400 rcf for 5 min at 4 °C. Cells were resuspended in 0.5% BSA in DPBS to 1,000 cells μl^−1^ concentration and immediately used for sequencing library preparation.

#### Flow cytometry of mouse splenocytes for quantification

Starting with single cell suspension, samples were aliquoted into new tubes so that final cell count was 1 × 10^7^ cells per sample, then centrifuged at 400 rcf 5 min at 4°C. Supernatant was removed, and samples were washed with 1mL PBS, then centrifuged again for 5 min. This was repeated 1x for a total of 2 PBS washes. Cells were then resuspended in 500μL of 1:500 UV Live/Dead Stain (ThermoFisher) in PBS and incubated at 4 °C for 30 min in the dark. Cells were washed with 500μL PBS, then centrifuged at 400 rcf 5 min at 4°C. Supernatant was removed, and the cell were resuspended in 1:50 Fc Block in MACS buffer so that the final concentration of cells was 1 × 10^7^ cells 100µL^−1^. Fluorophore-conjugated antibody cocktail was added in a 1:1 ratio. The following antibodies were used (antigen, clone, fluorophore, manufacturer, catalog no., dilution): I-A/I-E, 2G9, BUV395, BD, 743876, 1:1000; CD11b, M1/70, BUV737, BD, 612800, 1:500; F4/80, T45-2342, BV711, BD, 565612, 1:200; CD19, 1D3, BV786, BD, 563333, 1:500; CD3e, 145-2C11, BV786, BD, 564379, 1:200; NK1.1, PK136, BV786, BD, 740853, 3:1000; CD4, GK1.5, PE, Biolegend, 100407, 1:500; CD11c, HL3, PECF594, BD, 562454, 1:1000; CD63, NVG-2, PECy7, Biolegend, 143910, 1:1000; Tim-4, RMT4-54, AF647, BD, 564178, 1:10000; Tim-4, RMT4-54, PE, Biolegend, 130006, 1:10000; CD4, GK1.5, BUV395, BD, 563790, 1:500; CX3CR1, SA011F11, PE, Biolegend, 149006, 1:1000; CD4, RM4-5, eFluor450, BD, 50-112-8787, 1:1000; CD45.1, A20, BUV395, BD, 565212, 3:1000; CD45.2, 104, BV421, Biolegend, 109832, 3:1000; CD63, NVG-2, PECy7, Biolegend, 143910, 1:1000; CD44, IM7, BUV737, BD, 612799, 1:1000; CD45R/B220, RA3-6B2, BV605, Biolegend, 103243, 1:500; CD25, PC61, BV786, BD, 564023, 1:1000; CD4, GK1.5, Kiravia Blue 520, Biolegend, 100478, 1:500; CD3e, 145-2C11, PE, BD, 553064, 1:200; CD23, B3B4, PE Dazzle 594, Biolegend, 101634, 1:500; CD62L, MEL-14, PECy7, eBioscience, 25-0621-82, 1:1000; CD21/CD35, 7E9’, AF647, Biolegend, 123424, 1:500; CD8, 53-6.7, AF700, BD, 557959, 3:1000.

After incubation at 4 °C for 30 min in the dark, cells were washed with 750μl of MACS buffer and centrifuged at 400 rcf 5 min at 4°C. Supernatant was removed and cells were resuspended in 500μl 2% PFA in PBS/HBSS. Cells were incubated for 15 min at room temperature, then washed with 500μl of PBS. Cells were centrifuged, and supernatant was removed. Cells were resuspended in 500μl of PBS and stored in the dark at 4°C. Samples were run on the BD FACSymphony A5 Cell Analyzer. Compensation was performed using single-color controls and spectral unmixing, and weight optimization was performed to minimize spillover between primary channels.

#### Flow cytometry and cell sorting of mouse splenocytes for CITE-seq

Starting with single cell suspension, samples were aliquoted into new tubes so that final cell count was 2 - 5 × 10^7^ cells per sample, then centrifuged at 400 rcf 5 min at 4°C. Supernatant was removed, and the cells were resuspended in 1:50 Fc Block in PBS + 1% bovine serum albumin (BSA) so that the final concentration of cells was 1 × 10^7^ cells 100µL^−1^. Cells were incubated for 10 min at 4°C. Oligonucleotide-conjugated antibody cocktail was then added, and samples were mixed gently with wide bore pipette. Next, fluorophore-conjugated antibody cocktail was added. Final ratio of cells to cocktail (fluorophore-labeled plus oligonucleotide-labeled) was 1:1. Next, hashtag antibody was added to each sample at a ratio of 0.4 µL hashtag antibody per 1 × 10^7^ cells. Samples were incubated at 4C for 30 min in the dark. Samples were then washed with 4mL PBS + 1% BSA and centrifuged at 400 rcf for 5 min at 4°C. Supernatant was removed. This was repeated 3x for total of 4 washes. Samples were resuspended in PBS + 1% BSA to a concentration of 1 × 10^7^ cells ml^-1^. SYTOX Blue viability dye (Thermo Fisher) was added to cell suspension 5 minutes prior to sorting sample at a concentration of 1μl ml^−1^. If cell suspension contained visible clumps, sample was filtered using 30 µm pre-separation filters (Miltenyi Biotec) prior to sorting. Cells were sorted on a BD FACS Aria III SORP instrument using a 100-μm nozzle. Cells were sorted into 300μl of 2% BSA in Dulbecco’s phosphate-buffered saline (DPBS) and immediately pelleted by centrifugation at 400 rcf for 5 min at 4 °C. Cells were resuspended in 0.5% BSA in DPBS to 1,000 cells μl^−1^ concentration and immediately used for sequencing library preparation.

Oligonucleotide-labeled antibody cocktail was made fresh during day of experiment, to prevent clumping of antibodies. Antibodies were centrifuged at 14,000 rcf for 10 min prior to aliquoting antibody into cocktail, and care was taken to avoid pipetting from the bottom of the antibody vial, in order to avoid transferring any precipitated antibody present. TotalSeq-B Hashtag antibodies (BioLegend) were used for sample identification. TotalSeq-B antibodies (BioLegend) were used for protein quantification of selected markers. The following oligonucleotide-conjugated antibodies were used (antigen, clone, manufacturer, catalog no., final dilution): TotalSeq-B0301 Hashtag 1, M1/42; 30-F11, Biolegend, 155831, 1:500; TotalSeq-B0302 Hashtag 2, M1/42; 30-F11, Biolegend, 155833, 1:500; TotalSeq-B0303 Hashtag 3, M1/42; 30-F11, Biolegend, 155835, 1:500; TotalSeq-B0304 Hashtag 4, M1/42; 30-F11, Biolegend, 155837, 1:500; TotalSeq-B0114 F4/80, BM8, Biolegend, 123155, 1.6:1000; TotalSeq-B0001 CD4, RM4-5, Biolegend, 100573, 0.8:1000; TotalSeq-B0014 CD11b, M1/70, Biolegend, 101273, 1.6:1000; TotalSeq-B0106 CD11c, N418, Biolegend, 117359, 2.4:1000; TotalSeq-B0440 CD169, 3D6.112, Biolegend, 142429, 0.8:1000; TotalSeq-B0567 Tim4, RMT4-54, Biolegend, 130025, 0.08:1000; TotalSeq-B0563 Cx3cr1, SA011F11, Biolegend, 149045, 0.8:1000; TotalSeq-B0202 CD64, X54-5/7.1, Biolegend, 139329, 0.8:1000; TotalSeq-B0117 I-A/I-E, M5/114.15.2, Biolegend, 107657, 0.8:1000; TotalSeq-B0565 MERTK, 2B10C42, Biolegend, 151525, 0.8:1000.

For CITE-seq, fluorophore-conjugated antibody cocktail was made at twice the concentration used for standard flow cytometry. This ensured that the final antibody concentration during cell staining would be identical to what was used during standard flow cytometry. The following antibodies were used (antigen, clone, fluorophore, manufacturer, catalog no., final dilution): I-A/I-E, 2G9, BUV395, BD, 743876, 1:1000; F4/80, T45-2342, BV711, BD, 565612, 1:200; MERTK, DS5MMER, PE, eBioscience, 50-112-2328, 1:1000; Tim-4, RMT4-54, AF647, BD, 564178, 1:10000. Tim-4, RMT4-54, PE, Biolegend, 130006, 1:10000; CD63, NVG-2, PECy7, Biolegend, 143910, 1:1000; CD4, RM4-5, APC, eBioscience, 17-0042-81, 1:1000.

#### Cytospin analysis of mouse splenocytes

Starting with single cell suspension, samples were aliquoted into new tubes so that final cell count was 2 × 10^7^ cells per sample. Samples were washed with PBS, and then centrifuged at 400 rcf 5 min at 4°C. Supernatant was removed, and the cell were resuspended in 1:50 Fc Block in MACS buffer so that the final concentration of cells was 1 × 10^7^ cells 100µL^−1^. Fluorophore-conjugated antibody cocktail was added in a 1:1 ratio. The following antibodies were used (antigen, clone, fluorophore, manufacturer, catalog no., dilution): I-A/I-E, 2G9, BUV395, BD, 743876, 1:1000; F4/80, T45-2342, BV711, BD, 565612, 1:200; MERTK, DS5MMER, PE, eBioscience, 50-112-2328, 1:1000; Tim-4, RMT4-54, AF647, BD, 564178, 1:10000.

After incubation at 4 °C for 30 min in the dark, cells were washed with 750μl of MACS buffer and centrifuged at 400 rcf 5 min at 4°C. Samples were resuspended in MACS buffer to a concentration of 1 × 10^7^ cells ml^-1^ and taken to flow cytometry core on ice. SYTOX Blue viability dye (Thermo Fisher) was added to cell suspension 5 minutes prior to sorting sample at a concentration of 1μl ml^−1^, and samples were mixed gently prior to sorting. Cells were sorted on a BD FACS Aria III SORP instrument using a 100-μm nozzle. Cells were sorted into 300μl of 2% BSA in Dulbecco’s phosphate-buffered saline (DPBS) and immediately pelleted by centrifugation at 400 rcf for 5 min at 4 °C. Cytospin holders, funnels, filters, and slides were arranged appropriately. Cells were resuspended in MACS buffer, and 150ul aliquots were pipetted into cytospin funnels. Samples were centrifuged at 100 rcf for 5 minutes. Slides were detached carefully, then stained with Hema 3 Manual Staining System (Fisher HealthCare): slides were dipped into Fixative (methyl alcohol) x 5 seconds, Solution I (eosin) x 5 seconds, and Solution II (methylene blue) x 5 seconds. Slides were rinsed in tap water and then left to dry at room temperature. Cytoseal Mountant (Epredia) was used to mount coverslips prior to imaging. Brightfield 60x images of cytospin slides were taken on Olympus BX41 microscope equipped with an Olympus DP21 camera using Cellsens software, and final images were prepared using Fiji ImageJ software.

#### Recovery of cryopreserved human splenocytes

1 ml cell suspensions were warmed in 37°C water bath until just thawed, swirling gently to mix. Thawed cells were transferred into a 50 ml conical tube. RPMI 1640 without L-glutamine (Corning) with 5% FBS (Thermo-Fisher) and 200 μg/ml DNase (Roche) was used to dilute thawed sample. 50μl of the RPMI buffer was added every 5 seconds until sample was 5x original volume. Then, 100μl of the RPMI buffer was added every 5 seconds until sample was 10x original volume. Finally, 200μl of the RPMI buffer was added every 5 seconds until sample was 15x original volume. Sample was mixed gently via swirling throughout dilution process. If at this step, the cell suspension contained visible clumps, sample was then filtered using 30 µm pre-separation filters (Miltenyi Biotec). Sample was centrifuged at 400 rcf for 10 min at 4°C and resuspended in MACS buffer. Cell concentrations and viability were measured using a Cellometer K2 with AO/PI reagent. At this point, sample was ready for proceeding with protocol for human flow cytometry and cell sorting for scRNAseq.

#### Histology and immunohistochemistry

For histological analysis of human spleen, tissues were collected, rinsed in PBS, and excessive capsular tissue, if present, was removed prior to fixation. Human samples were fixed using 10% neutral buffered formalin (NBF) or 4% paraformaldehyde (PFA) in PBS for 48 hours at 4°C. Tissues were then stored in 70% EtOH. After a series of ethanol and xylene dehydration steps, spleen tissue was embedded in paraffin using Sakura Tissue-Tek VIP 5 instrument at 60°C at the Northwestern University Pathology Core Facility. Tissues were stained with either hematoxylin and eosin or antibodies for immunohistochemistry. For all immunohistochemistry analyses, heat-induced epitope retrieval was performed at pH 6.0 using citrate buffer. Then, slides were incubated in primary antibody overnight at 4°C. Slides were washed in PBS, and then stained with secondary HRP-conjugated antibody. Antibody binding was detected through incubation with DAB substrate until signal was seen, and reaction was then stopped. Haematoxylin counterstaining was performed.

Whole slide scanning of human tissue slides was performed using a Hamamatsu Nanozoomer 2.0-HT scanner, and quantification of the immunohistochemistry staining was done using QuPath^54^, Version v0.5.1-arm64. Briefly, the hematoxylin and eosin signal was estimated from the original RGB channel data using color deconvolution and estimated stain vectors derived from representative tissue regions. Then, representative tissue regions were selected from a subset of samples to use for training data. These regions were annotated manually and assigned tissue labels (“White pulp”, “Red pulp/vasculature”, and “Background/ignore”), which were then were used to train a random trees classifier using a very low resolution and the hematoxylin and eosin channels. The whole tissue sections were detected as regions to be analyzed using absolute thresholding in the red channel, and the outer 200 μm edge of tissue was excluded as background. Any tissue regions with large rips or similar significant disruption were excluded manually. Then, the random trees classifier was used to identify white pulp regions within the tissue sections, with manual annotation used to adjust the white pulp regions as needed. Next, absolute thresholding in the DAB channel was used to determine positive antibody signal. All tissues were analyzed using the same absolute thresholds, with no adjustment between samples. The createAnnotationsFromPixelClassifier Tool was used to generate annotations corresponding to positive antibody staining using the threshold, with a minimum size of 10 µm to exclude punctate positivity. This approach was used to quantify positive staining area in both the white pulp regions and the total tissue area. These areas were quantified out of total tissue area, and final measurements were exported into GraphPad for statistical testing and plot generation.

The following antibodies were used for immunohistochemistry (antigen, clone, manufacturer, catalog no., final dilution): DC-SIGN, DC28, Santa Cruz, sc-65740, 1:100; CD163, Polyclonal, Atlas Antibodies, HPA051974, 1:300; SIGLEC1, Polyclonal, Atlas Antibodies HPA053457, 1:250; TIMD4, Polyclonal, Atlas Antibodies, HPA015625, 1:1000; Human Osteoactivin/GPNMB, Polyclonal, R&D systems, AF2550, 1:500.

For H&E analysis of mouse spleen, spleens were collected and cut into 2-4 pieces. One piece per spleen was fixed for 48 hours at 4 °C. Samples for GM3-clo-lip experiments were fixed using 10% NBF; all other mouse samples were fixed using 4% PFA. After fixation, tissues were stored 70% EtOH. After a series of ethanol and xylene dehydration steps, spleen tissue was embedded in paraffin using Sakura Tissue-Tek VIP 5 instrument at 60°C at the Northwestern University Mouse Histology and Phenotyping Laboratory. Tissues were stained with hematoxylin and eosin. Whole slide scanning was performed using a Hamamatsu Nanozoomer 2.0-HT scanner, and quantification of white and red pulp regions was performed using QuPath^54^, Version v0.5.1-arm64. Briefly, the hematoxylin and eosin signal was estimated from the original RGB channel data using color deconvolution and estimated stain vectors derived from representative tissue regions. Then, representative tissue regions were selected from a subset of mice to use for training data. These regions were annotated manually and assigned tissue labels, which were then were used to train a random trees classifier using a very low resolution and the hematoxylin and eosin channels. The whole tissue sections were detected as regions to be analyzed using absolute thresholding in the red channel, and the outer 100 μm edge of tissue was excluded as background. Any tissue regions with large rips or similar significant disruption were excluded manually. Then, the random trees classifier was then run within the identified tissue sections, assigning labels to tissue regions. The area of these regions was quantified out of total tissue area, and final measurements were exported into GraphPad for statistical testing and plot generation.

#### Immunofluorescent microscopy

Mouse spleens were collected and cut into 2-4 pieces, depending on the experiment. One piece per spleen was taken for immunofluorescent microscopy and fixed for 3 hours in 2% PFA at 4°C, washed in PBS, then dehydrated in 20% sucrose (Sigma, S7903-1KG) overnight at 4°C. The next day, tissues were embedded in Tissue-Plus OCT compound (Fisher, 4585), flash frozen on dry ice, and placed in the –80°C. Using a cryotome, tissues were sectioned into 10 µm thick slices and placed immediately on microscopy slides (Fisherbrand Superforst Plus). Slides were dried at room temperature protected from light for 1 hour, then stored at 4°C until used for staining.

Prior to staining with antibody, tissues were rehydrated with PBS, and a barrier was drawn around individual tissue sections with a hydrophobic pen (Vector Laboratories). Tissues were then blocked and permeabilized in 0.2% Tween 20 (Biorad) in PBS with 10% Normal Donkey Serum (DS, Sigma) for 1 hour at room temperature. Next, tissues were stained with primary antibody in PBS-T with 10% DS overnight at 4°C. The next day, slides were washed 3x PBS-T, and slides were stained with secondary antibody in PBS-T with 10% DS at room temperature for 1 hour. Slides were washed 3x in PBS. Slides were then stained in Hoecshst (cat # 33342, invitrogen): 1:10,000 dilution in PBS for 5 min at toom temp. Finally, slides were washed once more with PBS. All of these steps were performed in a closed humidity chamber to keep tissues hydrated and protected from light. Finally, mounting solution (ProLong Diamond Antifade Mountant, ThermoFisher P36965) and a coverslip was then added to each slide one at a time. Slides were cured overnight in the dark at room temperature prior to imaging. After 24 hours, clear nail polish was used to seal the edges for long-term storage at −20°C.

Images were taken at 20x magnification using a Nikon Ti2B Widefield with Photometrix software. An LED light source and the following filters were used in the acquisition: Hoechst – Blue (Ex: 325-375 Em: 435-485), *ZsGreen* or *GFP* fluorescent reporter – Green (Ex: 450-490 Em: 500-550), MARCO – Red (Ex: 530-560 Em: 590-650), Tim-4 or CD169 – Far Red (Ex: 590-650 Em: 663-737). Quantification of images was done using QuPath^54^, Version v0.5.1-arm64. Briefly, 2-3 rectangular regions containing representative white pulp were manually identified in each mouse and annotated as regions of interest. Next, absolute thresholding in the Green, Red, and Far-Red channels was used to determine negative and positive signal for ZsGreen, MARCO, and CD169 signal, respectively. All tissues processed together were analyzed using the same absolute thresholds, with no adjustment between samples. The createAnnotationsFromPixelClassifier Tool was used to generate annotations corresponding to CD169+, MARCO +/- MMM and MARCO+, CD169-MZM using the established thresholds to set positive or negative staining, with a minimum size of 10 µm to exclude punctate positivity. Then, the ZsGreen threshold was used to determine ZsGreen positivity (no size exclusion). The percent area of the MMM and MZM annotations found to be ZsGreen-positive were quantified, and then compared in the untreated, 4-week, and 8-week timepoints. The data from each ROI was averaged within each mouse, and the average value from each mouse was treated an as independent sample when determining statistical significance between days. Final measurements were exported into GraphPad for statistical testing and plot generation.

The following primary antibodies were used for immunofluorescent micropscopy (antigen, clone, manufacturer, catalog no., final dilution): MARCO, EPR22944-64, abcam, ab239369, 1:500; CD169, 3D6.112, BioRad, MCA884GA, 1:500; Tim-4, RMT4-54, Biolegend, 130002, 1:250. The following secondary antibodies were used (species, target species, fluorophore, manufacturer, catalog no., final dilution): Donkey, anti-Rabbit IgG (H+L) Highly Cross-Adsorbed Secondary Antibody, Alexa Fluor 568. Invitrogen, A10042, 1:500; Donkey, anti-Rat IgG (H+L) Highly Cross-Adsorbed Secondary Antibody, Alexa Fluor Plus 647, Invitrogen, A48272, 1:500.

#### Single-cell RNA-seq

For human spleen samples, scRNA-seq was performed using Chromium Next GEM Single Cell 5′ V2 reagents (10x Genomics, protocol no. CG000331 Rev A). For mouse spleen samples, scRNA-seq was performed using the 10X Genomics 3’ V3.1 reagents (CG000317 Rev C), using a 10X Genomics Chromium X Controller. After quality checks, single-cell RNA-seq libraries were pooled and sequenced on Illumina NovaSeq 6000 or NextSeq 2000 instruments.

#### Analysis of scRNA-seq data

Data were processed using Cell Ranger 7.0.0 (10x Genomics) in exon-only mode, reads were mapped to GRCh38.98 (GENCODE v32/Ensembl98) or mm10 (GENCODE vM23/Ensembl98) reference genomes (10x Genomics Cell Ranger 2020-A package).

Mouse data was processed using Seurat V4^55^. For more details, see code. Human data were processed using Scanpy 1.7.2^56^, and multisample integration was performed with scvi-tools 0.14.0^57^. The scVI models was constructed on 1000 HVGs with the hyperparameters n_layers = 2, dropout_rate = 0.2, and n_latent = 10, and were trained using the settings max_epochs = 400, check_val_every_n_epoch = 2, and early_stopping = True. Default hyperparameters and settings were used otherwise. An initial round of Leiden clustering using the function sc.tl.leiden was performed on the integrated BAL object with a resolution of 0.75. Clusters characterized by low number of detected genes and transcripts and high percentage of mitochondrial genes were removed. Clusters containing doublets were identified as clusters simultaneously expressing lineage-specific marker genes (for example, C1QA for macrophages and CD3G for T cells) and excluded. Cell types were identified by marker genes, computed using the sc.tl.rank_genes_groups function with the settings method = “t-test”, n_genes = 200, and default settings otherwise. For more details, see code.

#### Single-cell spatial transcriptomics

FFPE mouse or human spleen tissue blocks were rehydrated in an ice bath for 30 min and sectioned on Leica HistoCore Biocut microtome at 5 um thickness. The first 5 sections were discarded, the tissue sections were floated in a 42°C water bath, mounted onto Xenium slides, and baked at 42°C for 3 hours, followed by deparaffinization, decrosslinking, probe hybridization, ligation, and amplification using Xenium In Situ Gene Expression workflow (CG000749_RevB). Samples were stained using Xenium Multi-Tissue Stain Mix and imaged on Xenium Analyzer instrument according to the manufacturer’s protocol (CG000749_RevB).

For analysis of the mouse spleen, we used a custom 363 gene panel (panel ID AREEUK; Supplemental Table 9). For analysis of the human spleen, we used Xenium Prime 5K Human Pan Tissue & Pathways Panel (10x Genomics) supplemented with 92 add-on genes (panel ID MVBJY8; Supplemental Table 10).

#### Analysis of mouse single-cell spatial transcriptomic data

Cell segmentation was performed with 10X Xenium onboard cell segmentation (Xenium Ranger v 2.0.0.10, 10x Genomics). For each sample, Xenium generated an output file of transcript information including x and y coordinates, corresponding gene target, assigned cell, a binary flag for whether the transcript was expressed over a nucleus, and quality score. Low-quality transcripts (qv < 20) and transcripts corresponding to blank/negative probes were removed.

Seurat v5 was used to perform further quality filtering and visualization. A single merged Seurat object was created for all samples based on ROI count matrices and metadata files with nuclei coordinates and area. Nuclei were retained according to the following criteria: ≥ 3 unique genes. Because Xenium outputs coordinates based on each slide, which results in samples with shared coordinates across multiple slides, the nuclei coordinates were manually adjusted for visualization so that no samples overlapped during plotting. These adjusted nucleus coordinates were added to the Seurat object and h5ad object as meta-data columns. Seurat v5 was used to perform dimensionality reduction, clustering, and visualization. Gene expression was normalized per cell using Seurat’s scTransform function. Dimensionality reduction was performed with PCA on 363_HVG. Cells were clustered based on this dimensionality reduction using the Louvain algorithm, using 30 PCs as input. Uniform Manifold Approximation and Projection (UMAP) plots of the data were generated using the same number of PCs as used for PCA. Cells were annotated using a combination of marker genes, generated using Seurat v5’s FindMarkers function, and spatial information, including cell morphology and position. The CellCharter package in Python was used to partition cells into 11 spatial niches using k-means clustering based on the transcriptomic expression of the nearest 6 neighboring cells. Spatial abundance analysis was performed by summing cells per cell type per sample. These per sample fractions were then compared across cell types using Students’s paired t test with FDR correction.

#### Analysis of human single-cell spatial transcriptomic data

Cell segmentation was performed with 10X Xenium onboard cell segmentation (Xenium Ranger v 2.0.0.10, 10x Genomics). For each sample, Xenium generated an output file of transcript information including x and y coordinates, corresponding gene target, assigned cell, a binary flag for whether the transcript was expressed over a nucleus, and quality score. Low-quality transcripts (qv < 20) and transcripts corresponding to blank/negative probes were removed.

Seurat v5 was used to perform further quality filtering and visualization. A single merged Seurat object was created for all samples based on ROI count matrices and metadata files with nuclei coordinates and area. Nuclei were retained according to the following criteria: ≥ 6 unique genes. Because Xenium outputs coordinates based on each slide, which results in samples with shared coordinates across multiple slides, the nuclei coordinates were manually adjusted for visualization so that no samples overlapped during plotting. These adjusted nucleus coordinates were added to the Seurat object and h5ad object as meta-data columns. Seurat v5 was used to perform dimensionality reduction, clustering, and visualization. Gene expression was normalized per cell using Seurat’s scTransform function. Dimensionality reduction was performed with PCA on 212 HVG. Cells were clustered based on this dimensionality reduction using the Louvain algorithm, using 30 PCs as input. Uniform Manifold Approximation and Projection (UMAP) plots of the data were generated using the same number of PCs as used for PCA. Cells were annotated using a combination of marker genes, generated using Seurat v5’s FindMarkers function, and spatial information, including cell morphology and position. The CellCharter package in Python was used to partition cells into 11 spatial niches using k-means clustering based on the transcriptomic expression of the nearest 6 neighboring cells. Spatial abundance analysis was performed by summing cells per cell type per sample. These per sample fractions were then compared across cell types using Students’s paired t test with FDR correction.

### Statistics and reproducibility

No statistical method was used to predetermine sample size. The experiments were not randomized. The Investigators were not blinded to allocation during experiments and outcome assessment. All statistics in the manuscript are reported as specified in the figure legends. When multiple hypothesis tests were performed, the false discovery rate (FDR) was controlled using the procedure of Benjamini and Hochberg. A significance level of 0.05 was used for all tests, unless indicated otherwise.

### Data availability

Single-cell RNA-seq counts tables and integrated objects are available through the Gene Expression Omnibus with accession numbers GSE326891. We also used publicly available single-cell RNA-seq data from GSE171124 and GSE223921. Xenium counts tables and Xenium Explorer bundles are available with accession number GSE326401. Single-cell and spatial data can be explored using interactive data browsers at: https://sqlifts.fsm.northwestern.edu/public/Thayer_2026/.

### Code availability

https://github.com/NUPulmonary/2026_Thayer.

## Acknowledgments

This research was supported in part through a generous gift from Kimberly Querrey and Louis A. Simpson. This research was supported by the Simpson Querrey Lung Institute for Translational Science (SQLIFTS), Northwestern University. This research was also supported by the computational resources and staff contributions provided for the Quest high-performance computing facility at Northwestern University, which is jointly supported by the Office of the Provost, the Office for Research and Northwestern University Information Technology. This research was also supported in part through the computational resources and staff contributions provided by the Genomics Compute Cluster, which is jointly supported by the Feinberg School of Medicine, the Center for Genetic Medicine and Feinberg’s Department of Biochemistry and Molecular Genetics, the Office of the Provost, the Office for Research and Northwestern Information Technology. The Genomics Compute Cluster is part of Quest, Northwestern University’s high-performance computing facility, with the purpose of advancing research in genomics. Northwestern University Flow Cytometry Core Facility is supported by the National Cancer Institute (NCI) Cancer Center support grant CA060553 awarded to the Robert H. Lurie Comprehensive Cancer Center. Cell sorting was performed on a BD FACS Aria SORP cell sorters purchased with the support of the National Institutes of Health (NIH, grant no. NIH 1S10OD011996-01 and 1S10OD026814-01). Integrative genomic services and single-cell spatial transcriptomics were performed by the Metabolomics Core Facility at Robert H. Lurie Comprehensive Cancer Center of Northwestern University. Next-generation sequencing was performed with support from the Simpson Querrey Institute for Epigenetics and at the Northwestern University NUSeq Core Facility. Imaging work was performed at the Northwestern University Center for Advanced Microscopy (RRID: SCR_020996), generously supported by NCI CA060553, awarded to the Robert H Lurie Comprehensive Cancer Center. Histology services were provided by the Northwestern University Mouse Histology and Phenotyping Laboratory which is supported by NCI CA060553 awarded to the Robert H Lurie Comprehensive Cancer Center. This work was supported by the Northwestern University Pathology Core Facility and a Cancer Center Support Grant (NCI CA060553). Spleen transplants were performed at the Microsurgery and Preclinical Research Core of the Comprehensive Transplant Center at Northwestern University. We acknowledge the use of tissues procured by the National Disease Research Interchange (NDRI; RRID:SCR_000550). We thank Dr. Nikolay S. Markov for advice and guidance on single-cell RNA-seq analysis. This work is dedicated to the benefit of all sentient beings.

R.A.G was supported by the Schmidt Science Fellows, in partnership with Rhodes Trust, and the Kimberly Querrey Fellowship in Data Science.

C.K. was supported by the NIH (grant no. HL176632).

A.B. was supported by the NIH (grant nos. P01HL169188, R01HL147290, R01HL145478, and R01HL147575).

H.P. was supported by the NIH (grant nos. R01AR080513, P01HL169188).

G.R.S.B. was supported by Simpson Querrey Lung Institute for Translational Science, the NIH (grant nos. P01AG049665, P01HL154998, U54AG079754, R01HL147575, R01HL158139, R01HL147290, and U19AI135964) and the Veterans Administration (award no. I01CX001777).

A.V.M. was supported by the NIH (grant nos. U19AI135964, P01AG049665, P01HL154998, P01HL169188, U19AI181102, R01HL153312, R01HL158139, and R01ES034350), and research grants from AbbVie and Merck.

## Author contributions

K.R.T. designed and executed mouse experiments, processed mouse and human tissues, performed FACS, cell sorting, scRNA-seq, IFM, analyzed data, performed computational and statistical analysis, and wrote the manuscript; M.J.S. performed computational analysis and data visualization and wrote the manuscript; Y.V.S. processed mouse and human tissues and performed IFM; R.A.N. assisted with mouse experiments, processed mouse and human tissues, processed tissues for Xenium analysis, and performed FACS and IFM; Z.Y. processed human tissues and performed scRNA-seq; Z.L. processed human tissues, performed scRNA-seq, and processed tissues for Xenium analysis; K.J.S. performed computational analysis; E.G.B. assisted with mouse experiments; W.T.P. assisted with mouse experiments; C.E.R. executed mouse experiments; R.A.G. executed mouse experiments; S.S. assisted with FACS; H.A-V. performed sequencing and spatial transcriptomics; C.K. provided human spleen samples; A.B. provided human spleen samples; A.D.Y. provided human spleen samples; R.P.M. provided human spleen samples; S.C.E. provided human spleen samples and assisted with Xenium data interpretation; N.S. F. provided human spleen samples and assisted with Xenium data interpretation; S.E.W. provided human spleen samples; M.C. assisted with regulatory and ethical oversight of the human studies; H.P. provided conceptual input and funding; G.R.S.B. provided conceptual input and funding, supervised the work and wrote manuscript; A.V.M. conceived, designed, and supervised the work, performed experiments and data analysis, and wrote the manuscript. All authors reviewed and approved the manuscript.

---

**Figure S1:**
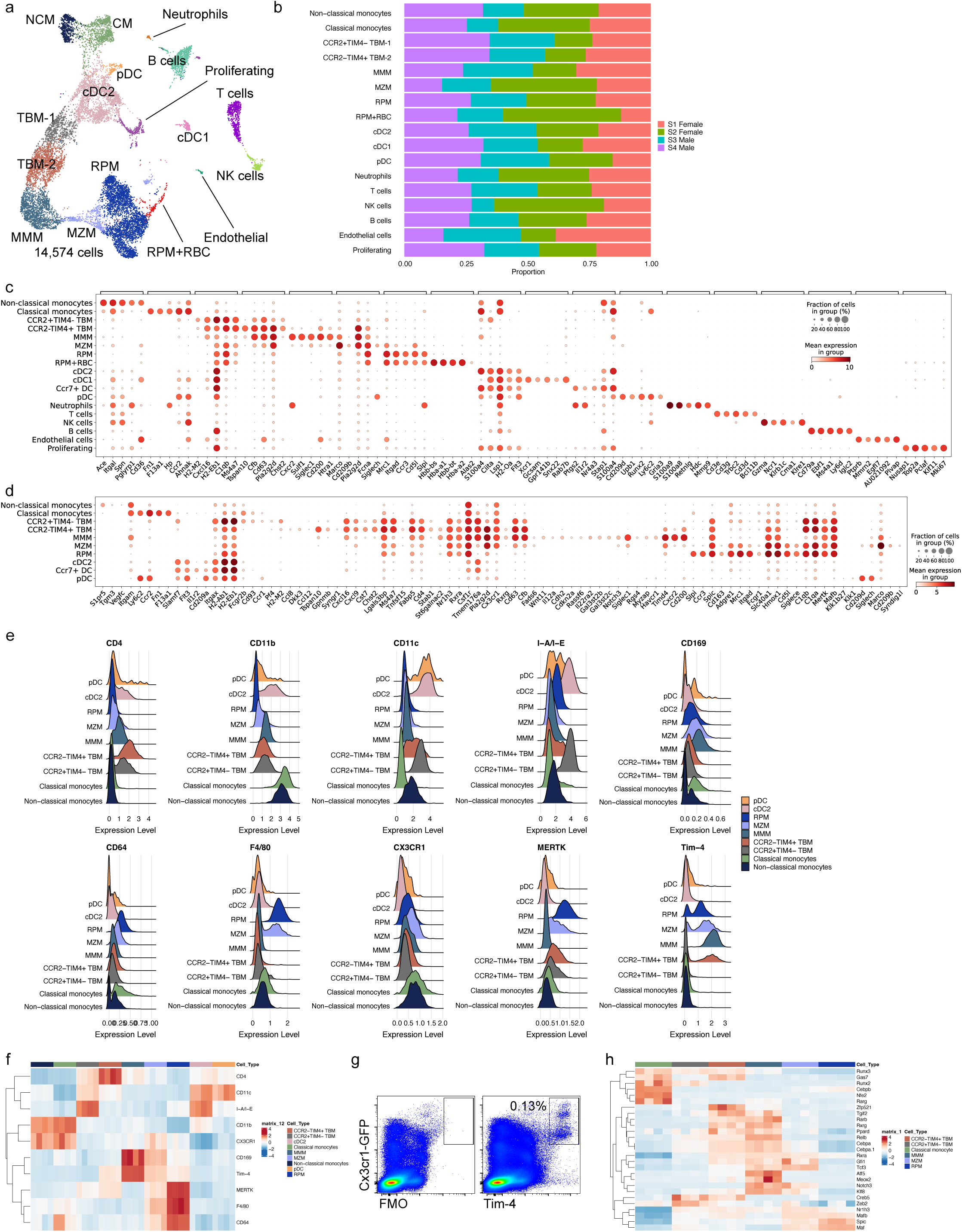
Identification and characterization of macrophages in mouse spleen. **a.** UMAP plot showing integrated analysis of 14,574 cells from 3–4-month-old *Cx3cr1^gfp/wt^* mice, 2 male and 2 female. b. Barplot illustrating composition of the clusters from UMAP on **a**. c. Dot plot illustrating expression of the top 5 cluster marker genes from **a**. d. Dot plot illustrating expression of selected genes within monocytes, macrophages, and dendritic cells from Figure 1d. Legend is the same as for panel **c**. e. Histograms illustrating the expression of selected proteins on monocytes, macrophages, and dendritic cells from Figure 1d as detected via the CITE-seq assay f. Heatmap illustrating the expression of selected proteins on monocytes, macrophages, and dendritic cells from Figure 1d as detected via the CITE-seq assay. Each column represents a single mouse. g. Cells with high GFP expression in *Cx3cr1^gfp/wt^* mice uniformly express high levels of Tim-4 protein detected via flow cytometry. Representative flow plots, gated on live singlets, showing fluorescence minus one control (FMO, left) and Tim-4 staining (right). Numbers indicate the percent of cells in the gate. h. Heatmap illustrating expression of selected transcription factors in monocytes and macrophages. Each column represents a single mouse.

**Figure S2:**
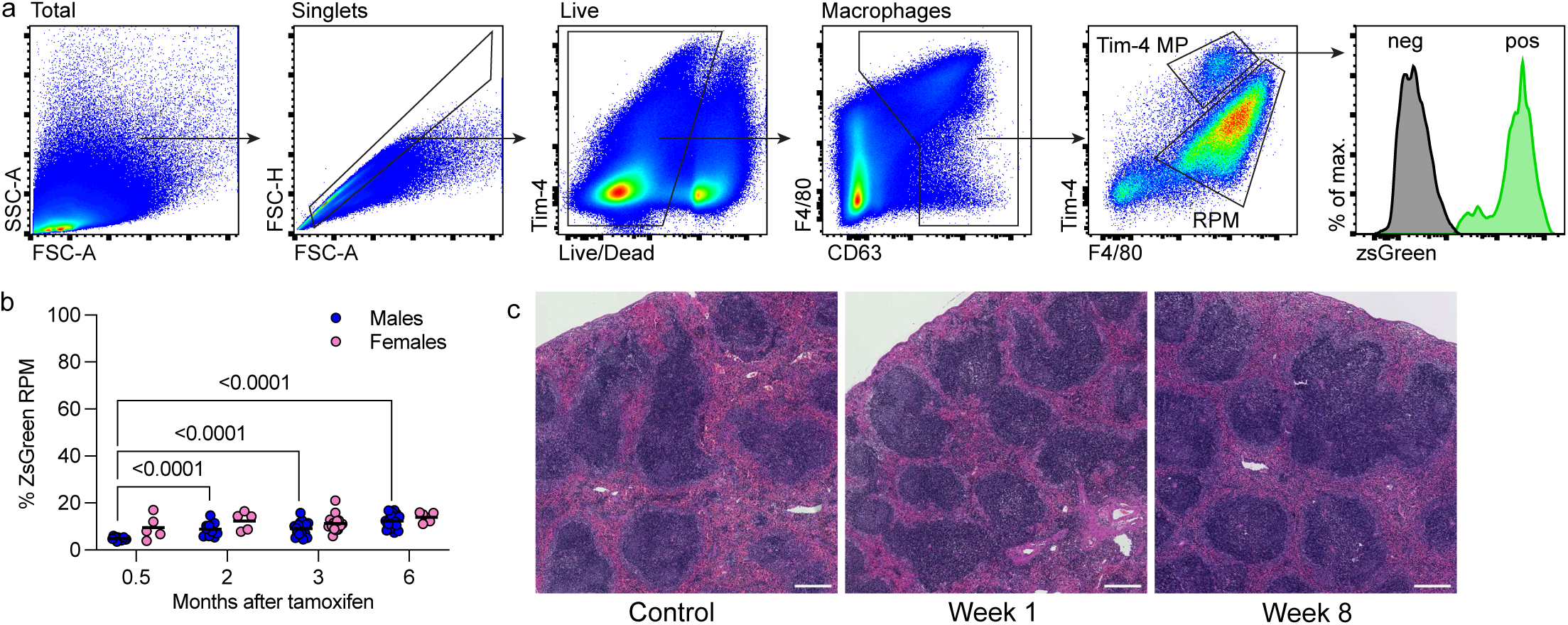
MMM and MZM are tissue-resident macrophages, MMM serve as a precursor population to MZM, and TBM have a dual ontogeny. **a.** Flow cytometry gating strategy for lineage-tracing experiments. Tim-4+ MP: Tim-4-positive macrophages. RPM: red pulp macrophages. b. Scatterplots showing percentage of ZsGreen-positive RPM in *Cx3cr1^ERCre^* x ZsGreen mice after tamoxifen treatment. Number of mice (from left to right): Female: 5, 5, 15, 6. Male: 13, 11, 17, 20. Results were compiled from 4 independent experiments. Significance was determined using one-way ANOVA, using a non-parametric Kruskal-Wallis test with Dunn’s multiple comparison test. The exact adjusted p-value is shown. c. Representative microphotographs of spleens from mice treated with GM3-CL. Scale bar is 250 μm.

**Figure S3:**
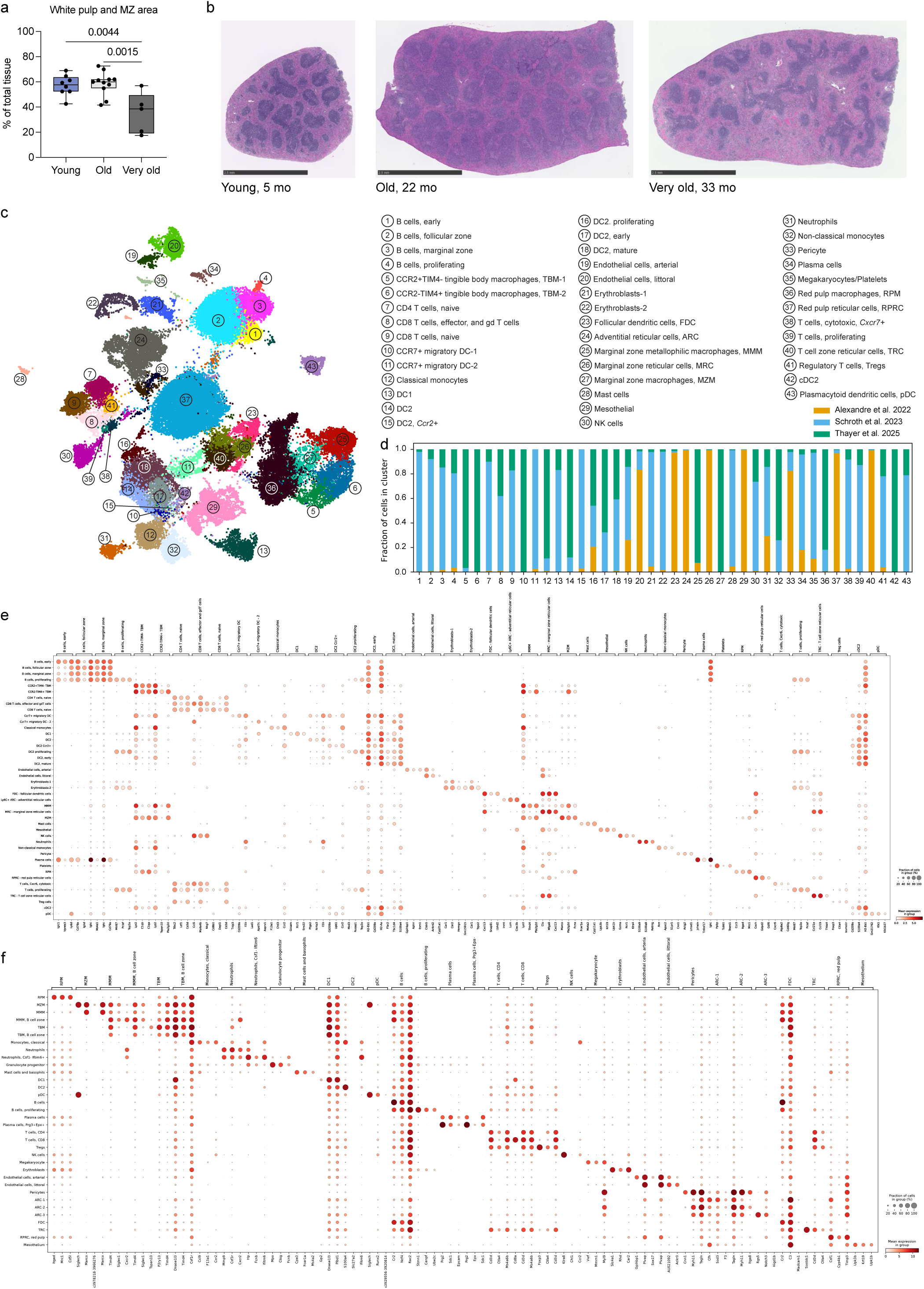
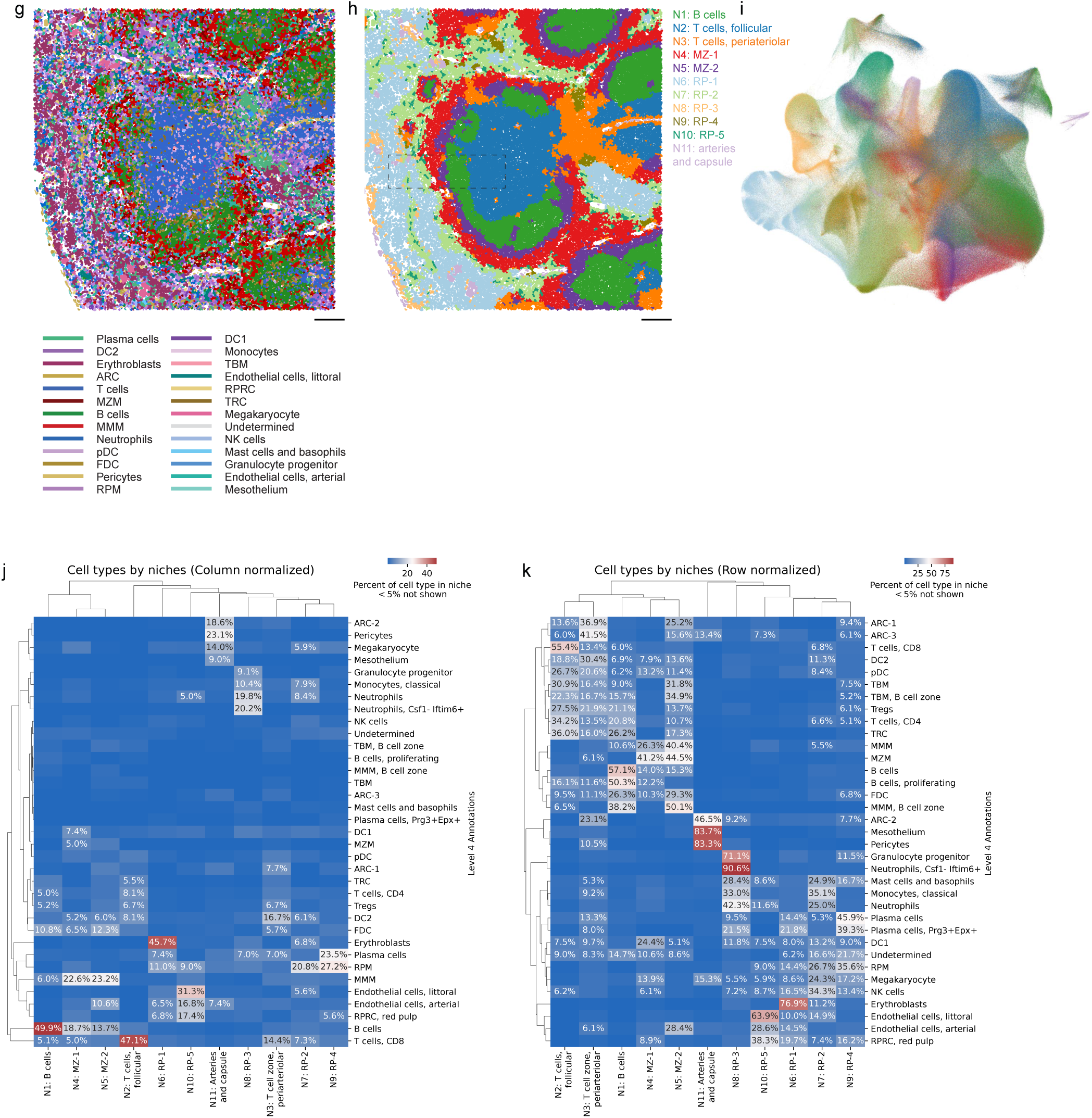
Changes in the aged mouse spleen are independent of a young circulation and include a decline in MMM and an increase in TBM, both of which remain tissue-resident cells. **a.** Box plot depicting quantification of white pulp and marginal zone area in young (n = 8, 3.5-5-month-old), old (n = 11, 22-month-old), and very old (n = 5, 27-33-month-old) C57BL/6 male mice. Quantification performed using QuPath software. One-way ANOVA followed by Tukey’s multiple comparison test was used to identify statistically significant differences between all groups. b. Representative H&E histology images illustrating age-related changes in splenic architecture. Scale bar is 2.5 mm. c. UMAP visualization of cell clusters from the integrated mouse single-cell spleen atlas. d. Bar plot illustrating cluster composition from mouse single-cell spleen atlas by study. e. Dot plot illustrating top 3 marker genes for each cluster from mouse single-cell spleen atlas. f. Dot plot illustrating top 3 marker genes for each cluster from mouse single-cell spatial spleen atlas. g. Spatial projection of cell types (level 3) resolved via spatial transcriptomics in the young mouse. Same spleen as in Figure 3b. The color palette is the same as Figures 3a**,b**. Scale bar is 100 μm. h. Spatial projection of niches resolved via spatial transcriptomics. Same spleen as in Figures 3b-d. Region matching Figures 3b-d is outlined with a dashed line. The color palette is the same as Figure 3d. Scale bar is 100 μm. i. Projection of the spatial niches onto UMAP from Figure 3a. The color palette is the same as Figure 3d. j. Heatmap illustrating the composition of spatial niches by cell type (level 4 annotations) in the integrated spatial object from Figure 3a (3 young, 4 old, and 4 very old mice). Hierarchical clustering on rows and columns, data normalized across columns. k. Heatmap illustrating the contribution of individual cell types (level 4 annotations) to spatial niches in the integrated spatial object from Figure 3a (3 young, 4 old, and 4 very old mice). Hierarchical clustering on rows and columns, data normalized across rows.

**Figure S4:**
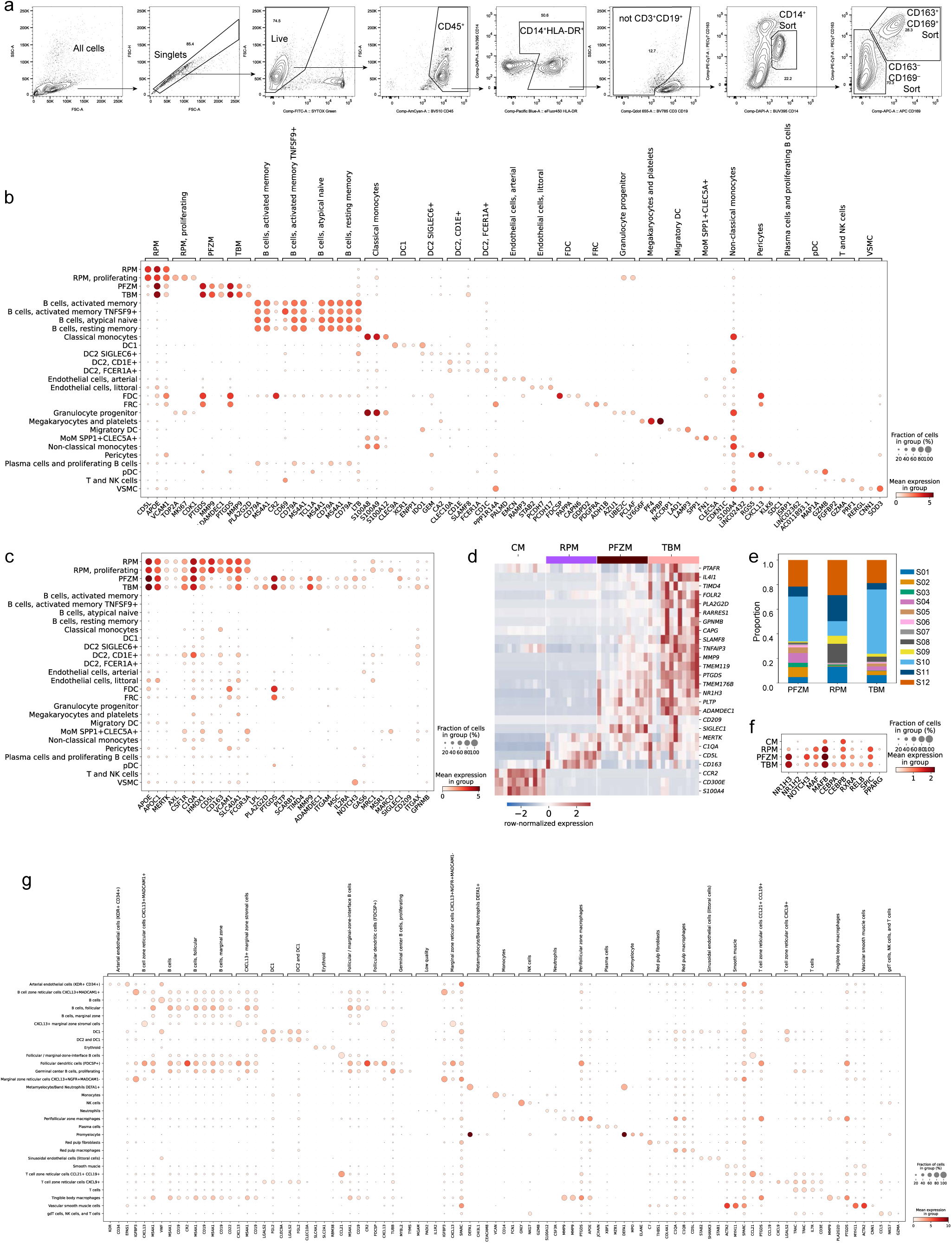
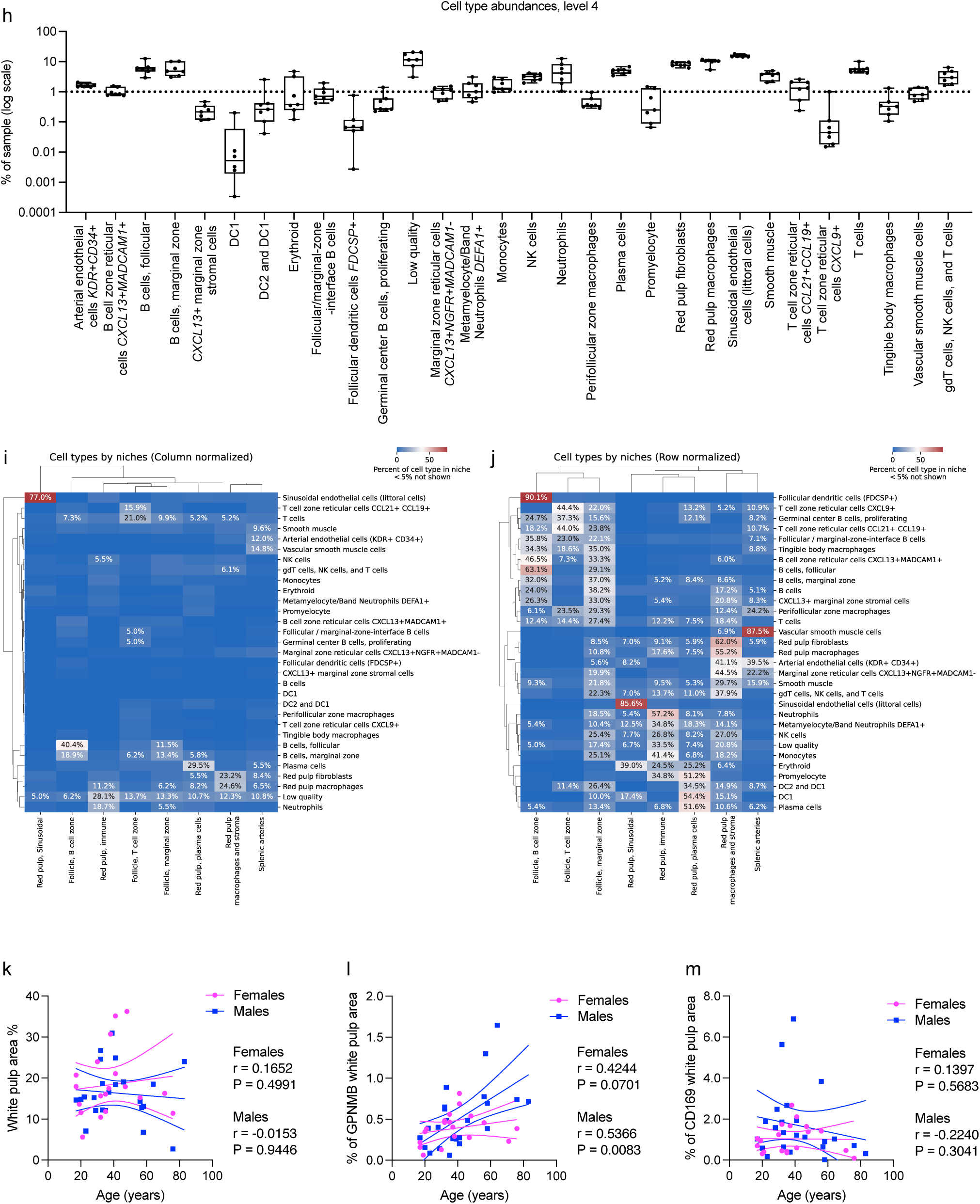
Identification of white pulp macrophages and their age-related changes in human spleen. **a.** Representative flow cytometry gating strategy for enriching macrophage subsets in the human spleen for scRNA-seq analysis. “Sort” indicates gates that were sorted and used for scRNA-seq analysis. b. Dot plot showing expression of top 3 marker genes in the integrated scRNA-seq analysis of 12 human spleens from Figure 4a. c. Dot plot showing expression of select genes in the integrated scRNA-seq analysis of 12 human spleens from Figure 4a. d. Heatmap showing expression of selected genes in tingible body macrophages (TBM), perifollicular zone macrophages (PFZM), red pulp macrophages (RPM), and classical monocytes (CM). e. Barplot showing composition of tingible body macrophages (TBM), perifollicular zone macrophages (PFZM), and red pulp macrophages (RPM). Every cluster is represented by cells from all 12 subjects. f. Dot plot showing expression of select transcriptional factors in tingible body macrophages (TBM), perifollicular zone macrophages (PFZM), red pulp macrophages (RPM), and classical monocytes (CM). g. Dot plot showing expression of top 3 marker genes in the integrated single-cell spatial transcriptomic analysis of 8 human spleens from Figure 4d. h. Box plot showing composition of the human spleen by cell type (level 4 annotations) using 7 high-quality spleen samples. Every cluster is represented by cells from all 7 subjects. i. Heatmap illustrating the composition of spatial niches by cell type (level 4 annotations) in the integrated spatial object from Figure 4d. Hierarchical clustering on rows and columns, data normalized across columns. Only 7 high-quality spleen samples out of 8 were included in the analysis. Numbers indicate the average across all 7 samples. j. Heatmap illustrating the contribution of individual cell types (level 4 annotations) to spatial niches in the integrated spatial object from Figure 4d. Hierarchical clustering on rows and columns, data normalized across rows. Only 7 high-quality spleen samples out of 8 were included in the analysis. Numbers indicate the average across all 7 samples. k. Scatter plot showing correlation between age and percentage of the splenic white pulp in spleens from 19 females and 23 males. The solid line illustrates the correlation, and the dashed line illustrates the 95% confidence interval. l. Scatter plot showing correlation between age and percentage of GPNMB-positive area within the splenic white pulp from 19 females and 23 males. The solid line illustrates the correlation, and the dashed line illustrates the 95% confidence interval. m. Scatter plot showing correlation between age and percentage of CD169-positive area within the splenic white pulp from 19 females and 23 males. The solid line illustrates the correlation, and the dashed line illustrates the 95% confidence interval.

## TABLES

Supplemental Table 1: Marker genes for the integrated mouse scRNA-seq object, related to Figure 1.

Supplemental Table 2: Marker genes for the integrated mouse scRNA-seq atlas, related to Figure 3.

Supplemental Table 3: Marker genes for the integrated mouse single-cell spatial transcriptomic object, related to Figure 3.

Supplemental Table 4: Hierarchical annotation of the integrated mouse single-cell spatial transcriptomic object, related to Figure 3.

Supplemental Table 5: Demographic characteristics of the human subjects, related to Figure 4.

Supplemental Table 6: Marker genes for the integrated human scRNA-seq object, related to Figure 4.

Supplemental Table 7: Marker genes for the integrated human single-cell spatial transcriptomic object, level 4 annotations, related to Figure 4.

Supplemental Table 8: Hierarchical annotation of the integrated human single-cell spatial transcriptomic object, related to Figure 4.

Supplemental Table 9: Xenium panel AREEUK. Supplemental Table 10: Xenium panel MVBJY8.

## REFERENCES

1. Lewis SM, Williams A, Eisenbarth SC. Structure and function of the immune system in the spleen. Sci Immunol [Internet]. American Association for the Advancement of Science (AAAS); 2019 Mar 1;4(33):eaau6085. Available from: 10.1126/sciimmunol.aau6085 PMCID: PMC6495537

2. Mebius RE, Kraal G. Structure and function of the spleen. Nat Rev Immunol [Internet]. Springer Science and Business Media LLC; 2005 Aug;5(8):606–616. Available from: 10.1038/nri1669 PMID: 16056254

3. den Haan JMM, Kraal G. Innate immune functions of macrophage subpopulations in the spleen. J Innate Immun [Internet]. S. Karger AG; 2012 Feb 7;4(5–6):437–445. Available from: 10.1159/000335216 PMCID: PMC6741446

4. Bronte V, Pittet MJ. The spleen in local and systemic regulation of immunity. Immunity [Internet]. Elsevier BV; 2013 Nov 14 [cited 2025 Feb 17];39(5):806–818. Available from: 10.1016/j.immuni.2013.10.010 PMCID: PMC3912742

5. Di Sabatino A, Carsetti R, Corazza GR. Post-splenectomy and hyposplenic states. Lancet [Internet]. 2011 July 2;378(9785):86–97. Available from: 10.1016/S0140-6736(10)61493-6 PMID: 21474172

6. Abila E, Buljan I, Zheng Y, Veres T, Weng Z, Nackenhorst M, Hulla W, Tolkach Y, Wöhrer A, Rendeiro AF. Tissue clocks derived from histological signatures of biological aging enable tissue-specific aging predictions from blood [Internet]. bioRxiv. 2024 [cited 2025 May 28]. p. 2024.11.14.618081. Available from: https://www.biorxiv.org/content/10.1101/2024.11.14.618081v1.abstract

7. Alex L, Rajan M, Xavier B, Jacob P, Rani K, Lakshmi G. Microscopic study of human spleen in different age groups. Int J Res Med Sci [Internet]. Medip Academy; 2015;1701–1706. Available from: 10.18203/2320-6012.ijrms2015055

8. Markus HS, Toghill PJ. Impaired splenic function in elderly people. Age Ageing [Internet]. Oxford University Press (OUP); 1991 July;20(4):287–290. Available from: 10.1093/ageing/20.4.287 PMID: 1927737

9. Ravaglia G, Forti P, Biagi F, Maioli F, Boschi F, Corazza GR. Splenic function in old age. Gerontology [Internet]. 1998;44(2):91–94. Available from: 10.1159/000021990 PMID: 9523220

10. Yona S, Kim KW, Wolf Y, Mildner A, Varol D, Breker M, Strauss-Ayali D, Viukov S, Guilliams M, Misharin A, Hume DA, Perlman H, Malissen B, Zelzer E, Jung S. Fate mapping reveals origins and dynamics of monocytes and tissue macrophages under homeostasis. Immunity [Internet]. 2013 Jan 24;38(1):79–91. Available from: 10.1016/j.immuni.2012.12.001 PMCID: PMC3908543

11. Kohyama M, Ise W, Edelson BT, Wilker PR, Hildner K, Mejia C, Frazier WA, Murphy TL, Murphy KM. Role for Spi-C in the development of red pulp macrophages and splenic iron homeostasis. Nature [Internet]. Springer Science and Business Media LLC; 2009 Jan 15;457(7227):318–321. Available from: 10.1038/nature07472 PMCID: PMC2756102

12. Haldar M, Kohyama M, So AYL, Kc W, Wu X, Briseño CG, Satpathy AT, Kretzer NM, Arase H, Rajasekaran NS, Wang L, Egawa T, Igarashi K, Baltimore D, Murphy TL, Murphy KM. Heme-mediated SPI-C induction promotes monocyte differentiation into iron-recycling macrophages. Cell [Internet]. 2014 Mar 13;156(6):1223–1234. Available from: 10.1016/j.cell.2014.01.069 PMCID: PMC4010949

13. Aichele P, Zinke J, Grode L, Schwendener RA, Kaufmann SHE, Seiler P. Macrophages of the splenic marginal zone are essential for trapping of blood-borne particulate antigen but dispensable for induction of specific T cell responses. J Immunol [Internet]. The American Association of Immunologists; 2003 Aug 1;171(3):1148–1155. Available from: 10.4049/jimmunol.171.3.1148 PMID: 12874200

14. Seiler P, Aichele P, Odermatt B, Hengartner H, Zinkernagel RM, Schwendener RA. Crucial role of marginal zone macrophages and marginal zone metallophils in the clearance of lymphocytic choriomeningitis virus infection. Eur J Immunol [Internet]. Wiley; 1997 Oct [cited 2025 Mar 24];27(10):2626–2633. Available from: https://pubmed.ncbi.nlm.nih.gov/9368619/ PMID: 9368619

15. Pirgova G, Chauveau A, MacLean AJ, Cyster JG, Arnon TI. Marginal zone SIGN-R1+ macrophages are essential for the maturation of germinal center B cells in the spleen. Proc Natl Acad Sci U S A [Internet]. Proceedings of the National Academy of Sciences; 2020 June 2 [cited 2025 Mar 24];117(22):12295–12305. Available from: https://pubmed.ncbi.nlm.nih.gov/32424104/ PMCID: PMC7275705

16. Baumann I, Kolowos W, Voll RE, Manger B, Gaipl U, Neuhuber WL, Kirchner T, Kalden JR, Herrmann M. Impaired uptake of apoptotic cells into tingible body macrophages in germinal centers of patients with systemic lupus erythematosus. Arthritis Rheum [Internet]. 2002 Jan;46(1):191–201. Available from: 10.1002/1529-0131(200201)46:1<191::AID-ART10027>3.0.CO;2-K PMID: 11817590

17. Schell SL, Soni C, Fasnacht MJ, Domeier PP, Cooper TK, Rahman ZSM. Mer receptor tyrosine kinase signaling prevents self-ligand sensing and aberrant selection in germinal centers. J Immunol [Internet]. Oxford University Press (OUP); 2017 Dec 15 [cited 2025 Mar 24];199(12):4001–4015. Available from: https://journals.aai.org/jimmunol/article-pdf/199/12/4001/1429743/ji1700611.pdf PMID: 29118245

18. Miyake Y, Asano K, Kaise H, Uemura M, Nakayama M, Tanaka M. Critical role of macrophages in the marginal zone in the suppression of immune responses to apoptotic cell-associated antigens. J Clin Invest [Internet]. American Society for Clinical Investigation; 2007 Aug [cited 2025 Mar 24];117(8):2268–2278. Available from: https://pubmed.ncbi.nlm.nih.gov/17657313/ PMCID: PMC1924497

19. Dijkstra CD, Döpp EA, Joling P, Kraal G. The heterogeneity of mononuclear phagocytes in lymphoid organs: distinct macrophage subpopulations in the rat recognized by monoclonal antibodies ED1, ED2 and ED3. Immunology [Internet]. Immunology; 1985 Mar [cited 2025 Mar 24];54(3):589–599. Available from: https://pubmed.ncbi.nlm.nih.gov/3882559/ PMCID: PMC1453512

20. Dijkstra CD, Van Vliet E, Döpp EA, van der Lelij AA, Kraal G. Marginal zone macrophages identified by a monoclonal antibody: characterization of immuno- and enzyme-histochemical properties and functional capacities. Immunology [Internet]. Immunology; 1985 May [cited 2025 Mar 24];55(1):23–30. Available from: https://pubmed.ncbi.nlm.nih.gov/3888828/ PMCID: PMC1453576

21. Kraal G, Janse M. Marginal metallophilic cells of the mouse spleen identified by a monoclonal antibody. Immunology [Internet]. Immunology; 1986 Aug [cited 2025 Mar 24];58(4):665–669. Available from: https://pubmed.ncbi.nlm.nih.gov/3733156/ PMCID: PMC1453104

22. Eikelenboom P. Characterization of non-lymphoid cells in the white pulp of the mouse spleen: an in vivo and in vitro study. Cell Tissue Res [Internet]. Springer Nature; 1978 Dec 29 [cited 2025 Mar 24];195(3):445–460. Available from: https://pubmed.ncbi.nlm.nih.gov/365349/ PMID: 365349

23. Humphrey JH. Tolerogenic or immunogenic activity of hapten-conjugated polysaccharides correlated with cellular localization. Eur J Immunol [Internet]. Wiley; 1981 Mar [cited 2025 Mar 24];11(3):212–220. Available from: https://pubmed.ncbi.nlm.nih.gov/7238567/ PMID: 7238567

24. Humphrey JH, Grennan D. Different macrophage populations distinguished by means of fluorescent polysaccharides. Recognition and properties of marginal-zone macrophages. Eur J Immunol [Internet]. Wiley; 1981 Mar [cited 2025 Mar 24];11(3):221–228. Available from: https://pubmed.ncbi.nlm.nih.gov/6940755/ PMID: 6940755

25. Jung S, Aliberti J, Graemmel P, Sunshine MJ, Kreutzberg GW, Sher A, Littman DR. Analysis of fractalkine receptor CX _3_ CR1 function by targeted deletion and green fluorescent protein reporter gene insertion. Mol Cell Biol [Internet]. Informa UK Limited; 2000 June;20(11):4106–4114. Available from: 10.1128/mcb.20.11.4106-4114.2000

26. Kraal G, Breel M, Janse M, Bruin G. Langerhans’ cells, veiled cells, and interdigitating cells in the mouse recognized by a monoclonal antibody. J Exp Med [Internet]. Rockefeller University Press; 1986 Apr 1;163(4):981–997. Available from: 10.1084/jem.163.4.981 PMCID: PMC2188075

27. Elomaa O, Kangas M, Sahlberg C, Tuukkanen J, Sormunen R, Liakka A, Thesleff I, Kraal G, Tryggvason K. Cloning of a novel bacteria-binding receptor structurally related to scavenger receptors and expressed in a subset of macrophages. Cell [Internet]. Elsevier BV; 1995 Feb 24;80(4):603–609. Available from: 10.1016/0092-8674(95)90514-6 PMID: 7867067

28. Gautier EL, Shay T, Miller J, Greter M, Jakubzick C, Ivanov S, Helft J, Chow A, Elpek KG, Gordonov S, Mazloom AR, Ma’ayan A, Chua WJ, Hansen TH, Turley SJ, Merad M, Randolph GJ, Immunological Genome Consortium. Gene-expression profiles and transcriptional regulatory pathways that underlie the identity and diversity of mouse tissue macrophages. Nat Immunol [Internet]. Springer Science and Business Media LLC; 2012 Nov;13(11):1118–1128. Available from: 10.1038/ni.2419 PMCID: PMC3558276

29. Orr SL, Le D, Long JM, Sobieszczuk P, Ma B, Tian H, Fang X, Paulson JC, Marth JD, Varki N. A phenotype survey of 36 mutant mouse strains with gene-targeted defects in glycosyltransferases or glycan-binding proteins. Glycobiology [Internet]. Oxford University Press (OUP); 2013 Mar;23(3):363–380. Available from: 10.1093/glycob/cws150 PMCID: PMC3605971

30. Chung JS, Dougherty I, Cruz PD Jr, Ariizumi K. Syndecan-4 mediates the coinhibitory function of DC-HIL on T cell activation. J Immunol [Internet]. The American Association of Immunologists; 2007 Nov 1;179(9):5778–5784. Available from: 10.4049/jimmunol.179.9.5778 PMID: 17947650

31. Umetsu SE, Lee WL, McIntire JJ, Downey L, Sanjanwala B, Akbari O, Berry GJ, Nagumo H, Freeman GJ, Umetsu DT, DeKruyff RH. TIM-1 induces T cell activation and inhibits the development of peripheral tolerance. Nat Immunol [Internet]. Springer Science and Business Media LLC; 2005 May 27 [cited 2025 Feb 23];6(5):447–454. Available from: https://www.nature.com/articles/ni1186 PMID: 15793575

32. Kobayashi N, Karisola P, Peña-Cruz V, Dorfman DM, Jinushi M, Umetsu SE, Butte MJ, Nagumo H, Chernova I, Zhu B, Sharpe AH, Ito S, Dranoff G, Kaplan GG, Casasnovas JM, Umetsu DT, Dekruyff RH, Freeman GJ. TIM-1 and TIM-4 glycoproteins bind phosphatidylserine and mediate uptake of apoptotic cells. Immunity [Internet]. Elsevier BV; 2007 Dec;27(6):927–940. Available from: 10.1016/j.immuni.2007.11.011 PMCID: PMC2757006

33. Nolte MA, Beliën JAM, Schadee-Eestermans I, Jansen W, Unger WWJ, van Rooijen N, Kraal G, Mebius RE. A conduit system distributes chemokines and small blood-borne molecules through the splenic white pulp. J Exp Med [Internet]. Rockefeller University Press; 2003 Aug 4;198(3):505–512. Available from: 10.1084/jem.20021801 PMCID: PMC2194088

34. Steiniger BS. Human spleen microanatomy: why mice do not suffice. Immunology [Internet]. 2015 July;145(3):334–346. Available from: 10.1111/imm.12469 PMCID: PMC4479533

35. Scott CL, T’Jonck W, Martens L, Todorov H, Sichien D, Soen B, Bonnardel J, De Prijck S, Vandamme N, Cannoodt R, Saelens W, Vanneste B, Toussaint W, De Bleser P, Takahashi N, Vandenabeele P, Henri S, Pridans C, Hume DA, Lambrecht BN, De Baetselier P, Milling SWF, Van Ginderachter JA, Malissen B, Berx G, Beschin A, Saeys Y, Guilliams M. The transcription factor ZEB2 is required to maintain the tissue-specific identities of macrophages. Immunity [Internet]. 2018 Aug 21;49(2):312–325.e5. Available from: 10.1016/j.immuni.2018.07.004 PMCID: PMC6104815

36. Okreglicka K, Iten I, Pohlmeier L, Onder L, Feng Q, Kurrer M, Ludewig B, Nielsen P, Schneider C, Kopfp M. PPARγ is essential for the development of bone marrow erythroblastic island macrophages and splenic red pulp macrophages. J Exp Med [Internet]. Rockefeller University Press; 2021 May 3;218(5). Available from: 10.1084/jem.20191314 PMCID: PMC8006858

37. A-Gonzalez N, Guillen JA, Gallardo G, Diaz M, de la Rosa JV, Hernandez IH, Casanova-Acebes M, Lopez F, Tabraue C, Beceiro S, Hong C, Lara PC, Andujar M, Arai S, Miyazaki T, Li S, Corbi AL, Tontonoz P, Hidalgo A, Castrillo A. The nuclear receptor LXRα controls the functional specialization of splenic macrophages. Nat Immunol [Internet]. Springer Science and Business Media LLC; 2013 Aug 16 [cited 2025 Feb 24];14(8):831–839. Available from: https://www.nature.com/articles/ni.2622 PMCID: PMC3720686

38. Wang Y, Zhang Y, Kim K, Han J, Okin D, Jiang Z, Yang L, Subramaniam A, Means TK, Nestlé FO, Fitzgerald KA, Randolph GJ, Lesser CF, Kagan JC, Mathis D, Benoist C. A pan-family screen of nuclear receptors in immunocytes reveals ligand-dependent inflammasome control. Immunity [Internet]. Elsevier BV; 2024 Dec 10;57(12):2737–2754.e12. Available from: 10.1016/j.immuni.2024.10.010 PMCID: PMC11634661

39. Fogg DK, Sibon C, Miled C, Jung S, Aucouturier P, Littman DR, Cumano A, Geissmann F. A clonogenic bone marrow progenitor specific for macrophages and dendritic cells. Science [Internet]. American Association for the Advancement of Science (AAAS); 2006 Jan 6;311(5757):83–87. Available from: 10.1126/science.1117729 PMID: 16322423

40. Madisen L, Zwingman TA, Sunkin SM, Oh SW, Zariwala HA, Gu H, Ng LL, Palmiter RD, Hawrylycz MJ, Jones AR, Lein ES, Zeng H. A robust and high-throughput Cre reporting and characterization system for the whole mouse brain. Nat Neurosci [Internet]. Springer Science and Business Media LLC; 2010 Jan;13(1):133–140. Available from: 10.1038/nn.2467 PMCID: PMC2840225

41. Grootveld AK, Kyaw W, Panova V, Lau AWY, Ashwin E, Seuzaret G, Dhenni R, Bhattacharyya ND, Khoo WH, Biro M, Mitra T, Meyer-Hermann M, Bertolino P, Tanaka M, Hume DA, Croucher PI, Brink R, Nguyen A, Bannard O, Phan TG. Apoptotic cell fragments locally activate tingible body macrophages in the germinal center. Cell [Internet]. 2023 Mar 16;186(6):1144–1161.e18. Available from: 10.1016/j.cell.2023.02.004 PMCID: PMC7614509

42. Gurwicz N, Stoler-Barak L, Schwan N, Bandyopadhyay A, Meyer-Hermann M, Shulman Z. Tingible body macrophages arise from lymph node-resident precursors and uptake B cells by dendrites. J Exp Med [Internet]. 2023 Apr 3;220(4). Available from: 10.1084/jem.20222173 PMID: 36705667

43. Nijen Twilhaar MK, Czentner L, Grabowska J, Affandi AJ, Lau CYJ, Olesek K, Kalay H, van Nostrum CF, van Kooyk Y, Storm G, den Haan JMM. Optimization of liposomes for antigen targeting to splenic CD169+ macrophages. Pharmaceutics [Internet]. MDPI AG; 2020 Nov 25;12(12):1138. Available from: 10.3390/pharmaceutics12121138 PMCID: PMC7760819

44. Birjandi SZ, Ippolito JA, Ramadorai AK, Witte PL. Alterations in marginal zone macrophages and marginal zone B cells in old mice. J Immunol [Internet]. Oxford University Press (OUP); 2011 Mar 15;186(6):3441–3451. Available from: 10.4049/jimmunol.1001271 PMCID: PMC3420341

45. Turner VM, Mabbott NA. Influence of ageing on the microarchitecture of the spleen and lymph nodes. Biogerontology [Internet]. 2017 Oct;18(5):723–738. Available from: 10.1007/s10522-017-9707-7 PMCID: PMC5597693

46. Schroth SL, Jones RTL, Thorp EB. Alloantigen Infusion Activates the Transcriptome of Type 2 Conventional Dendritic Cells. Immunohorizons [Internet]. 2023 Oct 1;7(10):683–693. Available from: 10.4049/immunohorizons.2300067 PMID: 37855737

47. Alexandre YO, Schienstock D, Lee HJ, Gandolfo LC, Williams CG, Devi S, Pal B, Groom JR, Cao W, Christo SN, Gordon CL, Starkey G, D’Costa R, Mackay LK, Haque A, Ludewig B, Belz GT, Mueller SN. A diverse fibroblastic stromal cell landscape in the spleen directs tissue homeostasis and immunity. Sci Immunol [Internet]. 2022 Jan 7;7(67):eabj0641. Available from: 10.1126/sciimmunol.abj0641 PMID: 34995096

48. Millard SM, Heng O, Opperman KS, Sehgal A, Irvine KM, Kaur S, Sandrock CJ, Wu AC, Magor GW, Batoon L, Perkins AC, Noll JE, Zannettino ACW, Sester DP, Levesque JP, Hume DA, Raggatt LJ, Summers KM, Pettit AR. Fragmentation of tissue-resident macrophages during isolation confounds analysis of single-cell preparations from mouse hematopoietic tissues. Cell Rep [Internet]. Elsevier BV; 2021 Nov 23;37(8):110058. Available from: 10.1016/j.celrep.2021.110058 PMID: 34818538

49. Gray EE, Friend S, Suzuki K, Phan TG, Cyster JG. Subcapsular sinus macrophage fragmentation and CD169+ bleb acquisition by closely associated IL-17-committed innate-like lymphocytes. PLoS One [Internet]. Public Library of Science (PLoS); 2012 June 1;7(6):e38258. Available from: 10.1371/journal.pone.0038258 PMCID: PMC3365896

50. Fujiyama S, Nakahashi-Oda C, Abe F, Wang Y, Sato K, Shibuya A. Identification and isolation of splenic tissue-resident macrophage sub-populations by flow cytometry. Int Immunol [Internet]. Oxford University Press (OUP); 2019 Feb 6;31(1):51–56. Available from: 10.1093/intimm/dxy064 PMCID: PMC6364618

51. Steiniger B, Bette M, Schwarzbach H. The open microcirculation in human spleens: a three-dimensional approach. J Histochem Cytochem [Internet]. SAGE Publications; 2011 June;59(6):639–648. Available from: 10.1369/0022155411408315 PMCID: PMC3201190

52. Steiniger BS, Seiler A, Lampp K, Wilhelmi V, Stachniss V. B lymphocyte compartments in the human splenic red pulp: capillary sheaths and periarteriolar regions. Histochem Cell Biol [Internet]. Springer Science and Business Media LLC; 2014 May [cited 2025 Dec 18];141(5):507–518. Available from: 10.1007/s00418-013-1172-z PMID: 24337546

53. Steiniger B, Barth P, Herbst B, Hartnell A, Crocker PR. The species-specific structure of microanatomical compartments in the human spleen: strongly sialoadhesin-positive macrophages occur in the perifollicular zone, but not in the marginal zone. Immunology [Internet]. Wiley; 1997 Oct;92(2):307–316. Available from: 10.1046/j.1365-2567.1997.00328.x PMCID: PMC1364073

54. Bankhead P, Loughrey MB, Fernández JA, Dombrowski Y, McArt DG, Dunne PD, McQuaid S, Gray RT, Murray LJ, Coleman HG, James JA, Salto-Tellez M, Hamilton PW. QuPath: Open source software for digital pathology image analysis. Sci Rep [Internet]. Nature Publishing Group; 2017 Dec 4;7(1):16878. Available from: 10.1038/s41598-017-17204-5 PMCID: PMC5715110

55. Hao Y, Hao S, Andersen-Nissen E, Mauck WM 3rd, Zheng S, Butler A, Lee MJ, Wilk AJ, Darby C, Zager M, Hoffman P, Stoeckius M, Papalexi E, Mimitou EP, Jain J, Srivastava A, Stuart T, Fleming LM, Yeung B, Rogers AJ, McElrath JM, Blish CA, Gottardo R, Smibert P, Satija R. Integrated analysis of multimodal single-cell data. Cell [Internet]. 2021 June 24;184(13):3573–3587.e29. Available from: 10.1016/j.cell.2021.04.048 PMCID: PMC8238499

56. Wolf FA, Angerer P, Theis FJ. SCANPY: large-scale single-cell gene expression data analysis. Genome Biol [Internet]. 2018 Feb 6;19(1):15. Available from: 10.1186/s13059-017-1382-0 PMCID: PMC5802054

57. Lopez R, Regier J, Cole MB, Jordan MI, Yosef N. Deep generative modeling for single-cell transcriptomics. Nat Methods [Internet]. 2018 Dec;15(12):1053–1058. Available from: 10.1038/s41592-018-0229-2 PMCID: PMC6289068

